# Arbovirus Vectors of Epidemiological Concern in the Americas: A Scoping Review of Entomological Studies on Zika, Dengue and Chikungunya Virus Vectors

**DOI:** 10.1101/713693

**Authors:** Reilly Jones, Manisha A. Kulkarni, Thomas M. V. Davidson, RADAM-LAC Research Team, Benoit Talbot

## Abstract

**Background:** Three arthorpod-borne viruses (arboviruses) causing human disease have been the focus of a large number of studies in the Americas since 2013 due to their global spread and epidemiological impacts: Zika, dengue, and chikungunya viruses. A large proportion of infections by these viruses are asymptomatic. However, all three viruses are associated with moderate to severe health consequences in a small proportion of cases. Two mosquito species, *Aedes aegypti* and *Aedes albopictus*, are among the world’s most prominent arboviral vectors, and are known primary vectors for all three viruses in the Americas.

**Objectives:** This review summarizes the state of the entomological literature surrounding the biology and ecology of vectors of Zika, dengue and chikungunya viruses and factors affecting virus transmission. The rationale of the review was to elucidate consensus and discord between studies, and guide future research based on identified knowledge gaps.

**Results:** A total of 196 studies were included in the scoping review after initial screening and subsequent exclusion of out-of-scope studies, secondary data publications, duplicate records, and studies unavailable in English language.

**Key findings:** Temperature and humidity have the strongest impact on mosquito distribution and dynamics, development of immatures and arborviral infection rates. Low socioeconomic status and related factors, including poor infrastructure, inconsistent access to water, and high household resident density, have been consistently associated with arbovirus vector occurrence. Effects of interspecific competition on arboviral vector species is currently poorly understood. Vector competence for Zika virus is well established for *Ae. aegypti* and *Ae. albopictus*. Information on Zika virus vector transmission dynamics is sparse in contrast to the wealth of research available for dengue and chikungunya viruses.

**Conclusions:** Based on the internationally recognized urgency of Zika virus infection as a public health concern, further research on arbovirus vectors and transmission dynamics is of pressing need.

## Introduction

Arboviruses are arthropod-borne viruses that comprise a diverse group of viruses transmitted largely by mosquitoes, including three viruses causing human disease with global implications: Zika, dengue, and chikungunya viruses. The term arbovirus does not encompass a taxonomically distinct group, but these viruses have similar life-history and transmission patterns that make information gleaned from one virus potentially useful to the understanding, and therefore prevention and control, of the others.

Since its identification in Zaire (now Democratic Republic of the Congo) in 1947, Zika virus (*Flavivirus, Flaviviridae*) has, until recently, largely been uncharacterized outside of Africa and Asia [1]. This virus generated international concern with its sudden emergence in Micronesia in 2007, followed by its introduction to French Polynesia in 2013 from somewhere in Southeast Asia [2]. The outbreak in French Polynesia affected more than 28,000 people. The virus ultimately reached the Americas in late 2014, resulting in the declaration of a Public Helath Emergency of International Concern by the World Health Organization [2]. To date, 86 countries have reported evidence of mosquito-transmitted Zika virus infection [3] Brazil currently faces the greatest burden of Zika virus infections [4]. Dengue fever, caused by four different serotypes of dengue virus (*Flavivirus, Flaviviridae*) is the most common arboviral disease that affects humans – 50 million people contract it each year, and an estimated 22,000 die from severe dengue [5]. Dengue is hyperendemic in the Americas, with cyclic epidemics occurring every three to five years [6]. Chikungunya virus (*Alphavirus, Togoviridae*) was first isolated in Tanzania in 1952 [7]. In the early 2000s, chikungunya virus cases and outbreaks were identified in countries in Africa, Asia, and Europe [7]. In 2013, it emerged in the Americas in Saint-Martin, and within the first year, over a million new cases were reported, spreading to 45 countries in the Latin American and Caribbean region [8].

A large proportion of Zika, dengue, and chikungunya viral infections are asymptomatic [9–11]. However, all three viruses are associated with moderate to severe health consequences in a small proportion of cases, with neonates, young children and/or older age groups at higher risk. Symptoms of Zika viral infection include rash, fever, arthralgia, and conjunctivitis [11]. More importantly, since its initial emergence in the Americas, Zika virus has been confirmed as a cause of congenital abnormalities (in infants born to women infected with Zika virus during pregnancy) and as a trigger of Guillain-Barré Syndrome [12]. Symptoms of dengue viral infection include rash, fever, arthralgia, and nausea. Some of the more severe symptoms of dengue viral infection may include deadly hemorrhage and plasma leak [9]. Symptoms of chikungunya viral infection include rash, fever, and arthralgia that may persist for months [7].

Two mosquito species, *Aedes aegypti* and *Aedes albopictus*, are among the world’s most prominent arboviral vectors. *Ae. aegypti* originated in Sub-Saharan Africa as a sylvatic species and was introduced to the Americas via ships soon after European arrival in the 1400s [13]. The species became domesticated and is now endemic to the Americas and the Asia-Pacific. The range of *Ae. albopictus* was restricted to Asia until the latter part of the 20th century. It is thought to have been introduced to the Western hemisphere through a shipment of used tires in 1985 and has expanded its territory to over 40% of the world’s landmass over the course of the past 30 years [14,15].

This review summarizes the state of the entomological literature surrounding the vectors of Zika, dengue and chikungunya viruses and factors affecting virus transmission. Waddell et al. [16] conducted a comprehensive scoping review of the Zika virus literature in 2016. However, the authors identified a limited scope of literature on vector studies, highlighting the need for a scoping review focusing on this area given its relevance in understanding arboviral disease risk. This scoping review aims to elucidate consensus and discord between studies, and guide future research based on identified knowledge gaps. Ultimately, the review was conducted to answer the following question: What is the current state of the evidence on mosquito vector population dynamics and behaviour, mosquito vector distribution and environmental suitability, vector species composition and virus transmission by mosquito vectors as they relate to Zika, dengue and chikungunya viruses in the Americas?

## Methods

This study’s search strategy and data extraction protocol were developed *a priori*. The list of definitions for each search term and the data characterization and utility form are available upon request. The preliminary search for this review was conducted on PubMed (National Library of Health, Bethesda, MD, United States). The search included the terms ‘zika’ OR ‘dengue’ OR ‘chikungunya’ AND ‘vector’ OR ‘Aedes aegypti’ OR ‘Aedes albopictus’. The search was conducted on March 1^st^ of 2018, and included all studies since January 1^st^ of 2013. Upon selection of potentially relevant articles, studies were characterized according to main characteristics including study setting, virus of interest, study design, methods of mosquito collection and analysis, vector species discussed, and main findings. Zotero (Center for History and New Media, George Mason University, United States) was initially used for title and abstract screening. All studies were subsequently transferred to Excel (Microsoft Corporation, Redmond, WA, United States) for data characterization and extraction. Two independent reviewers completed each step of the review following the broad initial screening, which was conducted by one reviewer.

### Inclusion Criteria

Articles were selected if they were related to mosquito population dynamics and behaviour, mosquito vector distribution and environmental suitability, mosquito vector species composition, and/or virus transmission by mosquito vectors, and if they were also related to Zika, dengue and/or chikungunya arboviruses. Other inclusion criteria included availability of an English language version, peer-review process, investigation of primary data, and for studies including a field component, specimen and data collection performed in the Americas. Studies that specifically examined the impacts of vector control measures, or that were unrelated to vector-borne aspects of disease, vector competence or entomological measures, were excluded due to the degree of scope expansion that would be caused by their inclusion.

## Results

### Descriptive Statistics of Scoping Review

The search yielded 6267 results. All records were screened, and 5919 were not deemed relevant based on title and abstract content. A total of 348 screened full-text studies were examined for eligibility, and ultimately 196 studies were included in the scoping review (Fig. 1; Table S1). The vast majority of studies were performed exclusively in the field, in the laboratory, or using a modelling framework, and were conducted exclusively on *Ae. aegypti*, or on *Ae. aegypti* and *Ae. albopictus* (Table 1). Studies focusing exclusively on dengue virus were the most numerous, while studies focusing exclusively on chikungunya virus were the least numerous (Table 1). Studies on mosquito population dynamics and behaviour were the most numerous, while studies on mosquito vector species composition or on multiple themes were the least numerous (Table 1). The average monthly number of studies hovered between 2 and 3 from 2013 to 2016, then increased to around 5 in 2017 and 2018 (Fig. 2), closely reflecting the introductions of Chikungunya and Zika viruses in the Americas and subsequent epidemics, respectively.

**Table 1.** Number of publications included in the scoping review, for each review section, study design, and arbovirus and mosquito vector species of interest.

| Theme | Category | Number of publications |
| --- | --- | --- |
| Section | Population Dynamics and Behavior | 54 |
|  | Spatial and Environmental Suitability | 46 |
|  | Vector Species Composition | 30 |
|  | Vector Transmission | 38 |
|  | Multiple | 28 |
| Study design | Field | 53 |
|  | Laboratory | 72 |
|  | Modelling | 44 |
|  | Field and Laboratory | 13 |
|  | Field and Modelling | 11 |
|  | Laboratory and Modelling | 3 |
| Virus of interest | Zika | 33 |
|  | Dengue | 76 |
|  | Chikungunya | 14 |
|  | Multiple | 38 |
|  | None specifically | 35 |
| Mosquito species of interest | *Ae. aegypti* | 105 |
|  | *Ae. albopictus* | 18 |
|  | *Cx. quinquefasciatus* | 4 |
|  | *Ae. aegypti* and *Ae. albopictus* | 38 |
|  | *Ae. aegypti* and *Cx. quinquefasciatus* | 2 |
|  | *Ae. albopictus* and *Cx. quinquefasciatus* | 1 |
|  | *Ae. aegypti*, *Ae. albopictus* and *Cx. quinquefasciatus* | 1 |
|  | Others | 25 |
|  | None specifically | 2 |

**Figure 1.**
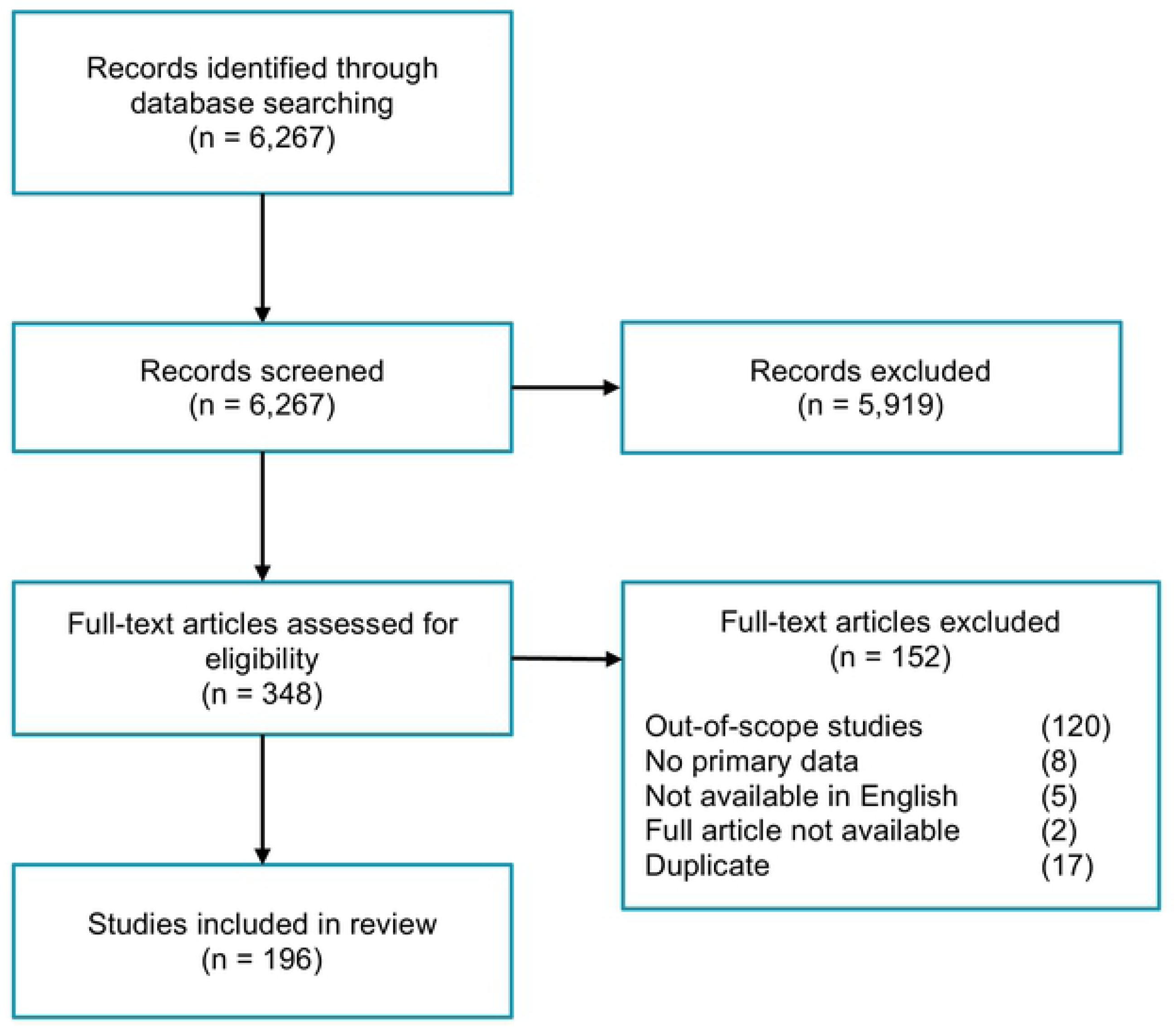
Summary of screening and exclusion steps of this scoping review’s methodology, and resulting number of publications after each step.

**Figure 2.**
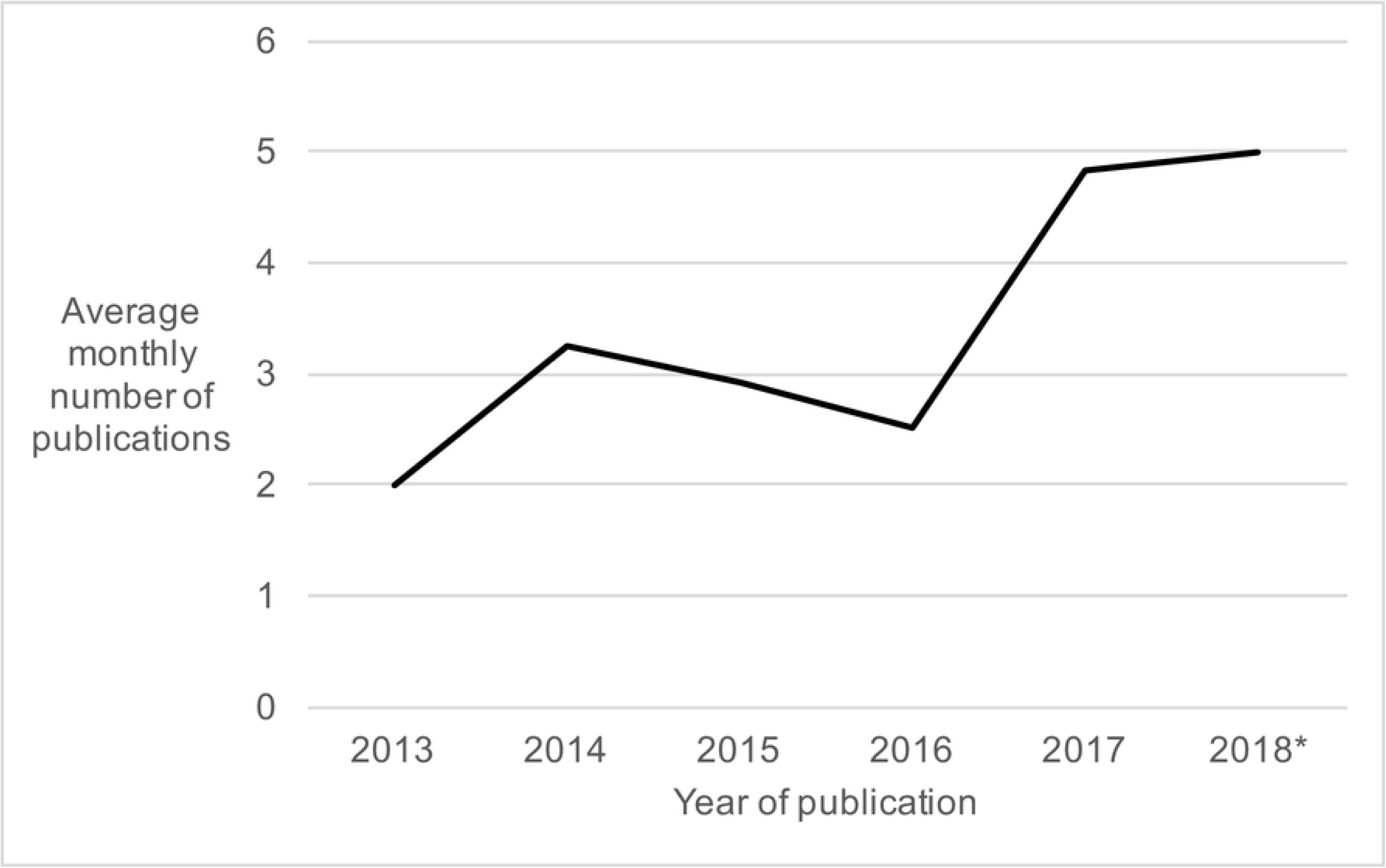
Average monthly number of publications included in the scoping review, for each year since 2013, out of a total of 196. *Year-to-date on March 1^st^ 2018.

### Population Dynamics and Behaviour

#### Meteorological and Seasonal Effects

A number of studies investigated the effects of meteorological factors on vector populations. Reiskind and Lounibos [17] conducted an observational field study in the United States in which they correlated meteorological variables and interspecific dynamics with mosquito crowding index. They found that the relative abundance of *Ae. aegypti* was higher than *Ae. albopictus* early in the wet season, and that interspecific mosquito crowding was highest one kilometer from the coast [17]. Degener et al. [18], Lucio et al. [19], Da Cruz Ferreira et al. [20] and Regis et al. [21] studied the impact of meteorological variables on *Ae. aegypti* infestation levels in South America. Degener et al. [18] found relative humidity to be the best predictor *Ae. aegypti* abundance, Lucio et al. [19] found saturated vapor pressure deficit to be most pertinent, and Da Cruz Ferreira [20] found mean temperatures above 18 °C and humidity levels below 75% to be positively predictive of *Ae. aegypti* abundance. As a secondary outcome, Degener et al. [18] found that MosquiTRAPs, which lure gravid *Ae. aegypti* through visual and olfactory stimuli, collected more *Ae. aegypti* during the dry season, and BG Sentinel traps, which mimic the convection currents of the human body, in addition to visual and olfactory stimuli, collected more during the rainy season [18].

Goindin et al. [22] experimentally studied the impact of temperature on female *Ae. aegypti* life history variables related to fecundity and life expectancy, and found that life expectancy varied with temperature, but fecundity did not. Costanzo et al. [23] conducted a similar experimental study with photoperiod as the independent variable, and found that *Ae. aegypti* and *Ae. albopictus* fencundity and life expectancy were higher in short photoperiod conditions.

Humidity was found to be highly important in statistical modelling analyses using trap capture data to model *Ae. aegypti* survival in Puerto Rico [24]. Padilla-Torres et al. [25] modelled *Ae. aegypti* and *Ae. albopictus* infestation rate according to meteorological variables, accounting for imperfect detection of vectors. Model-based infestation rate estimates were ∼30% higher than those calculated according to ovitrap data [25]. Two studies reported on the meteorological determinants of mosquito presence, and temporal population dynamics [26,27]. Sanavria et al. [26] found a strong correlation between temperature and *Ae. aegypti* presence in autumn (April, May and June) and a moderate correlation during the spring (October, November, December). Simoes et al. [27] found that *Ae. aegypti* populations tended to increase above a threshold of 18 °C and 54% humidity, and that extreme meteorological events were the best predictors of abundance.

#### Blood Feeding

Faraji et al. [28] conducted an observational field study in which they investigated geographical blood meal variation and blood meal source among *Ae. albopictus* and *Culex* species in the northeastern US. *Ae. albopictus* fed almost exclusively on mammalian hosts. More than 90% of their meals were derived from humans (58.2%) and domesticated pets (23.0% cats, 14.6% dogs). Vinauger et al. [29] experimentally demonstrated the importance of dopamine in relation to host-seeking behaviour.

#### Oviposition

Fader and Juliano [30] experimentally studied *Ae. albopictus* and *Ae. aegypti* oviposition site selection and found that both species preferred high detritus containers. Allgood and Yee [31] and conducted a similar experiment, and found that *Ae. albopictus* and *Cx. quinquefasciatus* prefer ovipositing in containers with organic chemical blends to those containing only water. Fonseca et al. [32] observed that *Ae. albopictus* preferred to oviposit in water with leaf infusion compared to water only, and that egg oviposition density was higher in the Fall than in the Summer [32]. Chitolina et al. [33] studied oviposition preference among *Ae. aegypti* between sewage water and clean water, and found that females had no preference for oviposition choice or and experienced no difference in oviposition success between sewage water and clean water. Williges et al. [34] and Rey and O’Connell [35] also studied oviposition site characteristic preferences. Williges et al. [34] studied oviposition height preference, and found that *Ae. albopictus* preferred to oviposit at ground level, compared to 1, 2, 3 or 4m in height under semi-field conditions. Rey and O’Connell [35] studied vegetation vs. structural component (wood and concrete blocks) oviposition site preference among *Ae. aegypti* and *Ae. albopictus*, and they also studied interspecific interactions. Interspecific interaction results were not conclusive, but *Ae. aegypti* were found to prefer vegetation for oviposition, and *Ae. albopictus* were found to prefer structural components [35]. Brown et al. [36] studied the combined effects of temperature and ecosystem type on distance travelled before oviposition, and found it to be limited at high temperature and low humidity. Santos et al. [37] studied the effects of WSMoL as an oviposition stimulant, and its mechanism. They found it to be effective under semi-field conditions through its effects on gustatory sensillia [37]. Wermelinger et al. [38], Valenca et al. [39] and Chadee and Martinez [40] observationally studied oviposition site preference of *Ae. aegypti*, and Cavalcanti et al. [41] used longitudinal modelling to do the same. Wermelinger et al. [38] found that gravid *Ae. aegypti* preferred to oviposit on the surface of the water in ovitraps. Valenca et al. [39] found laundry sinks to be the most common oviposition site type, while Chadee and Martinez [40] found holes in brick fences to be the most used. Cavalcanti [41] found ground tanks to be the most common oviposition site.

Estallo et al. [42] modelled the effects of meteorological variables on oviposition, and found minimum temperature (with a 3 week lag time) and weekly average rainfall to be the most important variables. De Majo et al. [43] similarly found that low temperature had a significant impact on *Ae. aegypti* egg survival, under experimental field conditions. Resende et al. [44] found moderate, positive correlation across collection measures from larval surveys, ovitraps and MosquiTRAPs. Dibo et al. [45] evaluated different egg collection techniques. They found that one sweep of the water’s surface was more suitable to collection in swimming pools, and the sequential five-sweep netting technique was more effective when used in drums and water-tanks [45]. Farnesi et al. [46] studied the effects of light exposure conditions on oviposition, and found that optimal viability requires a light/dark cycle, but oviposition in *Ae. aegypti* occurs preferentially in dark conditions.

Albeny-Simoes et al. [47] observed more frequent oviposition by *Ae. aegypti* after presence of predators in the same area, possibly in pursuit of conditions of greater bacterial abundance. In contrast, Wasserberg et al. [48] found that current nonlethal predator presence negatively affected *Ae. albopictus* oviposition behaviour. Davis et al. [49] found that *Ae. albopictus* was more likely to perform skip oviposition under crowded and poor nutrition conditions. Abreu et al. [50] also studied skip oviposition, and found that 7.4% of females exhibited skip oviposition, and that number of eggs oviposited was not affected by the density of available sites [50]. Wasserberg et al. [51] modelled the trade-off between intraspecific competition and oviposition site quality among *Ae. albopictus*, and Ruktanonchai et al. [52] studied *Ae. aegypti* behavioural plasticity of oviposition preferences under suboptimal conditions. Wasserberg et al. [51] found that oviposition activity first increased and then decreased with increasing intraspecific competition, in support of a hump-shaped model. Ruktanonchai et al. [52] found that previous experience with suboptimal oviposition conditions causes *Ae. aegypti* to modify their behaviour based on the assumption that nearby sites will also be suboptimal.

#### Larval Development

Gimenez et al. [53] experimentally studied container condition determinants of *Ae. aegypti* egg survival, and Byttebier et al. [54] studied the effects of growth medium and temperature among immatures. Giminez et al. [53] did not observe any difference in larval survival between conditions in which eggs were versus were not protected from macroscopic predators, under rising temperature conditions. Byttebier et al. [54] found hatching response (time to onset of hatching) to be lower at lower temperatures, and hatching rate to be higher on yeast media, compared to reverse osmosis water. Grech et al. [55] and Marinho et al. [56] found that development time varied inversely with temperature. De Majo et al. [57] found that egg hatching survival rate during the winter months in Brazil, was 30% at 13.2 °C and higher than 90% at 20 °C. They also observed development time was 49.9 days at 13.2 °C and 17.7 days at 20 °C [57]. Couret et al. [58] experimentally studied the effects of food availability, larval density and temperature on the development and survival of *Ae. aegypti*. Temperature was found to have the largest impact on juvenile mortality, and development was most variable at extreme values of diet and mosquito density. Lang et al. [59] experimentally studied the effect of larval food availability on adult *Ae. aegypti* male fitness and found it to be positively related to larger body size and greater swarming activity in adults. Lopes et al. [60] found no effect of isolation on survival and development of *Ae. aegypti* under typical seasonal conditions in Brazil. De Brito Arduino et al. [61] experimentally exposed *Ae. aegypti* to different salt concentrations, and found egg development occurred in all but the highest salt concentration (17.5%). Aznar et al. [62] modelled the growth rate at different development phases in *Ae. aegypti* and observed an impact of food restriction on growth during food dependent periods.

#### Competition

Noden et al. [63], Yee and Skiff [64], Riback et al. [65] examined interspecific and intraspecific competition on population dynamics. Noden et al. [63] found inter- and intraspecific competition to affect hatch-to-adult survivorship and development time to adulthood in *Ae. aegypti* and *Ae. albopictus*, and Riback et al. [65] made similar observations. Yee and Skiff [64] found *Ae. albopictus* female mass to be negatively affected by the presence of *Cx. coronator*. Alto et al. [66] studied the impact of *Ae. aegypti* and *Ae. albopictus* competition on female adult survival under high and low humidity. Intraspecific but not interspecific competition was found to reduce adult survival under both conditions. Wagman et al. [14] experimentally studied the range expansion and interspecific dynamics of *Ae. aegypti* and *Ae. albopictus* in Belize, and found *Ae. aegypti* was more abundant than *Ae. albopictus*. Kesavaraju et al. [67] studied the interspecific dynamics of *Ae. albopictus* and *Ae. sirrensis* in the United States and found *Ae. sirrensis* population growth decreased with increasing *Ae. albopictus* population density.

#### Population Genetics

Two studies examined *Ae. aegypti* population genetics, as they relate to their spatial distribution and genetic clustering [68,69]. The study conducted in Colombia by Jaimes-Duenez et al. [68] reports low genetic differentiation over time and two distinct genetic groups, which they inferred came from independent introductions. The study conducted by Steffler et al. [69] in Brazil reports high genetic diversity within populations, and attributes the lack of population differentiation partially to human motorized travel. Kotsakiozi et al. [70] report a genetic break between northern and southern Brazilian *Ae. aegypti* populations.

### Vector Distribution and Environmental Suitability

#### Microenvironment

Two studies reported the occurrence of *Ae. aegypti* infestation on marine vessels [71,72]. Guagliardo et al. [71] found mosquito abundance on barges to be highly aggregated, and Guagliardo et al. [72] found large barges to account for the majority of observed *Ae. aegypti* abundance (71.9%). Five studies examined the occurrence and correlates of domiciliary and peri-domicilliary mosquito density, including sociodemographic characteristics, container types, and meteorological correlates [73–77]. Specifically, Rodrigues et al. [73] and Serpa et al. [77] studied *Aedes aegypti* and *Aedes albopictus*, and Favaro et al. [74], Morales-Pérez et al. [75] and Schafrick et al. [76] studied *Aedes aegypti*. Covered containers [75–77], low education of the household head [75] and higher densities of residents per household [73] were found to be associated with higher risk of infestation. Two studies reported storm drain characteristics associated with mosquitoe presence: one related to *Aedes aegypti* and *Culex quinquefasciatus* presence [78], and the other to *Aedes aegypti* and *Aedes albopictus* [79]. Another reported the density of mosquitoes in storm drains compared to other settings [80]. Overgaard et al. [81] and Unlu et al. [82] investigated container characteristics associated with the presence of particular mosquito species. One study evaluated the correlation between mosquito resting habitat and energy accumulation in Florida, US [83]. *Ae. aegypti* was the most commonly collected species, was most often found on Mexican petunia, and their lipid and glycogen storage was different depending on vegetation type on which they were found. Quintero et al. [84] reported *Ae. aegypti* occurrence according to container type across five countries, and found vector population growth was favoured when containers were uncovered, outdoors, unused, and rain-filled in the dry season. Yee et al. [85] performed cluster analyses according to general microenvironmental characteristics.

#### Socioecological Factors

Wilke et al. [86] examined determinants of *Aedes aegypti, Aedes albopictus, Aedes fluviatilis, Aedes scapularis, Culex nigripalpus* and *Culex quinquefasciatus* dynamics in urban parks in São Paulo, Brazil. They reported on the impact of urban development as well as monthly temperature and accumulated rainfall over the month of collection and the previous month on species richness and abundance in that context [86]. They found temperature to significantly impact species abundance, but that correlation differed by species [86]. Montagner et al. [87] found *Ae. albopictus* to be more common in urban environments compared to rural, and similar results were obtained for *Ae. aegypti* by Overgaard et al. [81], Hagenlocher et al. [88] and Silva Lima et al. [89]. Fuentes-Vallejo et al. [90] compared *Ae. aegypti* density between two municipalities, and found high density to be associated with unplanned urbanization, flood-prone areas, low socioeconomic strata, household water tanks, high temperature, and lack of control measures for adult mosquitoes.

One study measured the utility of the premises condition index (PCI) as a predictor of *Ae. aegypti* infestation in Merida, Mexico [91]. PCI incorporates considerations of tidiness, patio shade, and house maintenance. Tidiness and shade were found to be useful dimensions, and houses with lower PCI scores (higher house maintenance, higher tidiness and lower shading of the patio) had significantly lower *Ae. aegypti* infestation and productivity [91]. Espinosa et al. [92] used vector breeding, density and cluster data to produce GIS models of *Ae. aegypti* spatial patterns in Argentina. Proportion of urban coverage, urban form, and water supply most strongly predicted *Ae. aegypti* distribution [92]. In a similar modelling study, Little et al. [93] found that decaying infrastructure and vegetation levels significantly influenced *Ae. albopictus* distribution. Miller and Loaiza [15] studied determinants of the expansion of *Ae. albopictus* in Panama, and found that the road network best predicted distribution over time. Taber et al. [94] modelled the colonization of Pennsylvania by *Ae. albopictus* as well as corresponding risk of dengue, and found that junkyard land cover and mean temperatures around 26 °C were associated with the highest risk of infestation.

#### Climatic Suitability

Eight studies examined arbovirus vector distribution and ecological suitability in the Americas. Five were conducted in the United States [93,95–98], two in Brazil [99,100], and one in Peru [101]. Johnson et al. [97] created a predictive model of population dynamics in the United States based on vector presence data, Little et al. [93] found that *Ae. albopictus* spatial variability was greater than annual temporal variability in the United States, and Carvalho et al. [100] found that *Ae. albopictus* was present in 59% of analyzed Brazilian municipalities. Across these studies, we can conclude that mosquitoes are often highly clustered in their distribution [99,101] and that distributional determinants may differ between *Ae. aegypti* and *Ae. albopictus* [96,97,100]. Seven studies predicted the distribution of arbovirus vectors under conditions of climate change [102–108]. One was conducted in Brazil [102], one in Ecuador [103], one in Mexico [104], one in Argentina [105], one in the United States [108], one in relation to the North American continent [106], and one in relation to the Northern Hemisphere [107]. Fischer et al. [105] predicted higher *Ae. aegypti* peak abundances and oviposition season duration in Argentina under conditions of climate change. Cardoso-Leite et al. [102] report that 80% of Brazil’s population currently resides in *Ae. aegypti* suitable regions, and that total risk area will decrease and spread southwards by 2050. Studies related to the Northern Hemisphere predict increases in climatically suitable regions [106–108]. One study retrospectively studied the presence of *Ae. aegypti* and *Ae. albopictus* in the United States between 1995-2016 [109]. Two studies made predictions about vector range related to a specific disease under climate change – one related to dengue in the United States [110] and one related to chikungunya in Canada [111].

#### Mosquito Species Composition

Five studies examined mosquito species composition at the neighbourhood level [112–116]. Three were conducted in Brazil [113–115], one in Haiti [112] and one in Argentina [116]. Both large-scale studies conducted in Brazilian parks in Sao Paulo found *Ae. albopictus* to be the most abundant species [113,114]. Another study conducted in Brazil led to the collection of a small number of mosquito larvae in artificial and natural breeding sites, all of which were identified as *Ae. aegypti* [115]. The study conducted in Haiti found that an earthquake made conditions more suitable to *Ae. aegypti* infestation [112]. The study conducted in Argentina led to the collection of mosquitoes in twenty-seven cities, and all collections included either *Culex* spp. or *Ae. aegypti*. Cities with a mean annual temperature above 14.5°C were all found to be positive for *Ae. aegypti* [116].

### Vector Species Composition

#### Zika Virus

There is extensive evidence that *Ae. aegypti* mosquitoes are able to transmit Zika virus in both the laboratory [117–128] and in the field [129,130]. *Ae. albopictus* mosquitoes were also able to transmit Zika virus in experimental studies [120,122], but studies in which both *Ae. aegypti* and *Ae. albopictus* were captured found no Zika virus-infected *Ae. albopictus* [129,130]. Gendernalik et al. [131] and O’Donnell et al. [124] report that *Ae. vexans* mosquitoes are also experimentally competent vectors of Zika virus, but no studies indicated natural *Ae. vexans* infection with Zika virus.

*Cx. quinquefasciatus* has been identified by predictive models as a potential vector for Zika virus [132], as have *Sabethes* and *Haemagogus spp*. [133]. Seven studies found that *Cx. quinquefasciatus* mosquitoes were refractory to Zika virus when exposed to infectious blood meals [128,134–139]. Ferreira-de-Brito et al. [129] reported that no *Cx. quinquefasciatus* captured in Brazil were positive for Zika virus. Also, Guedes et al. [140] detected Zika virus in the midgut, salivary glands and saliva of artificially fed *Cx. quinquefasciatus* captured in Brazil, using RT-PCR and transmission electron microscopy. The same study also reported Zika virus isolated from two field-caught *Cx. quinquefasciatus* in Brazil. Aliota et al. [141] report that *Cx. pipiens* is refractory to Zika virus.

#### Dengue Virus

*Ae. albopictus* [126,142–145] and *Ae. aegypti* [126,143,144,146–148] are both experimentally competent to transmit dengue virus. Infection by the virus is observed in field populations of *Ae. albopictus* [149–152], *Ae. aegypti* [149,150,152–160] and *Cx. quinquefasciatus* [154], although the latter was not identified as a competent vector species experimentally.

#### Chikungunya Virus

*Ae. aegypti* [126,127,144,161–164,164–166], *Ae. albopictus* [127,144,162,164–167], *Aedes terrens* [168], and *Haemagogus leucocelaenus* [168] are all experimentally competent to transmit chikungunya virus. Chikungunya virus transmission in *Ae. aegypti* has also been observed in the field [119,121,157,169,170].

### Vector Transmission

#### Vector Competence Factors

Four studies measured the effect of temperature on vector competence [145,161,162,171]. Adelman et al. [161] found that under silenced RNAi conditions, *Ae. aegypti* were more predisposed to chikungunya infection at lower temperatures. Alto et al. [162] found that larger fluctuations in diurnal temperature range led to higher rates of chikungunya infection, and Xiao et al. [145] found that maximum dengue infection rates occurred at 31 °C. Mordecai et al. [171] modelled *Ae. aegypti* and *Ae. albopictus* transmission in the Americas and found that mean temperature data accurately reflected Zika, chikungunya and dengue human case data. Transmission was found to occur between 18 and 34°C and maximal transmission was observed between 26-29°C, with less certainty surrounding the critical thermal minimum than the critical thermal maximum [171]. *Ae. albopictus* was found to perform better in cooler temperatures [171]. Buckner et al. [143] found that the interaction of low temperature and low food availability increased *Ae. aegypti* and *Ae. albopictus* susceptibility to DENV-1 serotype infection.

Three studies examined the effects of larval competition on dengue vector competence [142,143,172]. Bara et al. [142] found that *Ae. albopictus* larval competition resulted in significantly longer development times, lower emergence rates, and smaller adults, but did not significantly affect the extrinsic incubation period of DENV-2 virus. Kang et al. [172] found that larval-stage crowding and nutritional limitation led to lower survival rates until pupation, lower blood feeding success, slower development, smaller adult body size, and lower susceptibility to DENV-2 infection. Four studies examined a variety of blood meal characteristics on arboviral infection rate [122,123,147,173]. Fresh Zika-infected blood meal was associated with significantly higher infection rates than frozen Zika-infected blood meal [122]. Similarly, Zika-infected whole blood meal was associated with significantly higher infection rates than Zika-infected protein meal [123]. Hill et al. [147] studied the impact of antibiotics on dengue infection rate and mosquito fertility, and found no significant association in *Ae. aegypti*. Mosquitoes exposed to DENV-2 were more likely to re-feed than those that were unexposed [173]. Sylvestre et al. [174] studied the impact of DENV-2 infection on *Ae. aegypti* life history traits, and found that it significantly affected feeding behaviour, survival, fecundity, and oviposition success.

#### Vector Infection Rate

Two studies conducted in Brazil exclusively examined infection rates by Zika virus in wild mosquito populations (Table 2). Ferreira-de-Brito et al. [129] reported three Zika-infected pools of *Ae. aegypti*, but no Zika-infected *Cx. quinquefasciatus* or *Ae. albopictus* pool [129], out of 468 tested pools among the three species. Ayllón et al. [130] tested 406 *Ae. aegypti* and 11 *Ae. albopictus* field-collected individuals, and found three Zika-infected *Ae. aegypti* individuals.

**Table 2.** List of studies that report a proportion of positive mosquito pools for any or a combination of Zika, dengue and chikungunya viruses, along with information on authors, year and country of location of the study, and mosquito species of interest.

| Authors | Year | Location | Mosquito species | Pools tested | Zika rate (%) | Dengue rate (%) | Chikungunya rate (%) |
| --- | --- | --- | --- | --- | --- | --- | --- |
| Ferreira-de-Brito et al. | 2016 | Brazil | *Aedes* sp. and *Cx. quinquefasciatus* | 468 | 0.64 | ø | ø |
| Ayllon et al. | 2017 | Brazil | *Ae. aegypti* and *Ae. albopictus* | 178 | 1.12 | ø | ø |
| Martinez et al. | 2014 | Mexico | *Ae. aegypti* | 226 | ø | 0.88 | ø |
| Calderon-Arguedas et al. | 2015 | Costa Rica | *Ae. albopictus* | 35 | ø | 25.71 | ø |
| Cecilio et al. | 2015 | Brazil | *Aedes* sp. | 54 | ø | 7.41 | ø |
| Cruz et al. | 2015 | Brazil | *Ae. aegypti* | 50 | ø | 16.00 | ø |
| Perez-Castro et al. | 2016 | Colombia | *Ae. aegypti* | 34 | ø | 61.76 | ø |
| Perez-Perez et al. | 2017 | Colombia | *Ae. aegypti* and *Ae. albopictus* | 407 | ø | 32.43 | ø |
| Diaz-Gonzalez et al. | 2015 | Mexico | *Ae. aegypti* | 557 | ø | ø | 3.23 |
| Cevallos et al. | 2018 | Ecuador | *Ae. aegypti* | 22 | 14.29 | ø | 12.50 |
| Dzul-Manzanilla et al. | 2015 | Mexico | *Ae. aegypti* | 284 | ø | 9.51 | 3.17 |
| Cigarroa-Toledo et al. | 2016 | Mexico | *Ae. aegypti* | 27 – 237* | ø | 0.00 | 0.84 – 7.40* |
| Farraudiere et al. | 2017 | Martinique | *Ae. aegypti* | 414 | ø | 1.21 | 2.66 |

\*Total number of pools tested is not stated, but number of sampled mosquitoes, and maximum number of mosquitoes per pool, are stated.

Six studies reported exclusively on dengue infection rates in wild mosquito populations (Table 2). Cecílio et al. [175] observed four positive pools, out of 54 tested, among *Aedes* mosquitoes collected in two regions of Brazil over the course of 17 months, through the installation of ovitraps in public schools. Cruz et al. [155] detected eight positive pools, out of 50 *Ae. aegypti* pools, collected in Mato Grosso, Brazil. Martínez et al. [160] reported two positive pools, out of 226 *Ae. aegypti* pools, collected in Mexico. Claderón-Arguedas et al. [176] reported nine positive pools, out of 35 *Ae. albopictus* pools, collected in Costa Rica. Pérez-Pérez et al. [152] reported 132 positive pools, out of 407 tested, collected in Colombia. One of the positive pools was *Ae. albopictus*, and the remainder were *Ae. aegypti*. Pérez-Castro et al. [177] reported 21 positive pools, out of 34 tested, in *Ae. aegypti* in Colombia.

A study measured the naturally-occurring prevalence of chikungunya virus in wild mosquito populations (Table 2). Díaz-González et al. [170] reported 18 *Ae. aegypti* positive pools in Mexico, out of 557 tested. A study reported on the prevalence of both chikungunya and Zika viruses among *Ae. aegypti* in Ecuador (Table 2) [121]. Three studies tested both chikungunya and dengue viruses in wild mosquito populations (Table 2). Chikungunya, but not dengue, was detected in *Ae. aegypti* in Mexico by Cigarroa-Toledo et al. [169], although both chikungunya and dengue viruses were isolated in Mexico in *Ae. aegypti* by Dzul-Manzanilla et al. [157], and in Martinique by Farraudière et al. [159].

#### Vertical Transmission

Three studies reported on vertical transmission of dengue virus [156,158,178], and one [179] reported on the vertical transmission of Zika virus. Buckner et al. [178] found a vertical transmission rate of DENV-1 of 11.11% in *Ae. albopictus* and of 8.33% in *Ae. aegypti*. Da Costa et al. [156] observed dengue infection rates among third and fourth instar *Ae. aegypti* between 1.14% and 2.41% among Brazilian municipalities, and Espinosa et al. [158] observed one DENV-3 positive male *Ae. aegypti* pool, collected in Argentina. Thangamani et al. [179] experimentally injected mosquitoes with Zika virus and observed Zika virus infection in *Ae. aegypti* offspring, but not *Ae. albopictus*. Six filial *Ae. aegypti* pools out of 69 tested were found positive for Zika virus [179].

#### Transmission Risk Modelling

Five studies modelled transmission dynamics for Zika virus [138,180–183]. Lourenço et al. [138] used vectorial capacity as a means of prediction, Marini et al. [180] and Majumder et al. [185] used vector abundance and human case data, and Villela et al. [182] and Ospina et al. [183] used disease notification and natural history.

Eight studies modelled dengue transmission dynamics [184–191]. Lee et al. [192] constructed a predictive model that accurately foresaw 75% of dengue outbreaks in Colombia. Reiner et al. [184] reported that social proximity drives fine-scale heterogeneity in dengue transmission rates based on data from Peru. Rojas et al. [185] found attack rates in Girardot and San Andres, Colombia to be highest among females, aged 20-49. Three studies reported that meteorological variables including temperature and humidity are important determinants of transmission dynamics [186,187,189,190], and one study found that transovarial transmission plays an important role in transmission dynamics depending on basic reproductive number [188]. Liu-Helmersson et al. [107] predicted an increase in diurnal temperature range and increased dengue epidemic potential under climate changes in cold, temperate and extremely hot climates where mean temperatures are far from 29 °C. Fitzgibbon et al. [193] and Velasques-Castro et al. [194] studied *Ae. aegypti* dynamics in relation to host spatial heterogeneity. Fitzgibbon et al. [193] report that early host and vector heterogeneity significantly affect final epidemic size, and Velasquez-Castro et al. [194] generated a dengue infection risk map, based on host dynamics.

One study estimated chikungunya transmission risk according to temperature threshold for breeding and adult mosquitoes in Argentina [195]. The authors suggest that temperatures conducive to *Ae. aegypti* breeding and transmission are present during September and April in northeastern Argentina, and in January in southern Argentina. A study compared endemic and transient chikungunya and dengue transmission dynamics, and the role of virus evolution [196]. They found that reducing biting rate and vector-to-susceptible-host ratio were the most effective at reducing basic reproductive number. A study modelled transmission risk of Zika, dengue and chikungunya and found temperature data to match well with human case data [171].

#### Strain Infectivity and Co-Infection

Six studies examined the infectivity of different dengue viral strains, and the impact of co-infection [148,172,197–200]. Quintero-Gil et al. [198] found that the DENV-2 serotype performed with a thousand-fold greater efficiency than the DENV-3 serotype, upon co-infection. Vazeille et al. [200] found that DENV-4 outperformed DENV-1 in *Ae. aegypti* upon co-infection. Muturi et al. [148] found that infection with DENV-4 rendered *Ae. aegypti* significantly less susceptible to secondary infection with DENV-2. In parallel, Serrato-Salas et al. [199] found that *Ae. aegypti* were significantly less susceptible to secondary dengue infection, after having been challenged with an inactive version of the virus. Kang et al. [172] modelled interactions between dengue viral serotypes. Quiner et al. [197] studied the infectivity of different isolates of DENV-2, and found NI-2B to have a replicative advantage over NI-1 until 12 days following infection, after which the advantage had dissipated. Nuckols et al. [144] artificially infected *Ae. aegypti* and *Ae. albopictus* with chikungunya and DENV-2 simultaneously, separately, and in reverse order. Simultaneous dissemination was detected in all groups upon co-infection, and co-transmission occurred at low rates [144]. Ruckert et al. [126] found that the co-infection of *Ae. aegypti* with Zika, chikungunya and dengue viruses minimally affected vector competence, and that vectors were able to transmit each viral pair, as well as three viruses simultaneously. Alto et al. [167] found *Ae. aegypti* and *Ae. albopictus* to be susceptible to Indian Ocean and Asian chikungunya virus genotypes.

#### Human Disease Risk

Five articles studied correlations between entomological measures and risk of human dengue infection [201–205]. One study conducted in Peru found that *Ae. aegypti* density was not associated with an increased risk of seroconversion [201]. One study in Acre, Brazil found that *Ae. aegypti* density and risk of dengue increased with tourism and case importation [202]. A study in Mexico City found a positive correlation between dengue incidence and *Ae. aegypti* indoor abundance, as well as monthly average temperature and rainfall [203]. Another study conducted in Peru found that an individual’s likelihood of being bitten in the home was directly proportional to time spent in the home, and body surface area. They did not find age and gender to be significant predictors [204]. Oliveira et al. [205] reported the circulation of four dengue serotypes in Brazil introduced between 2001 and 2012 (DENV-1, DENV-2, DENV-3, DENV-4) and reported an increase in dengue infection in Brazil during that time period, i.e. 587 cases/100 000 in 2001 to 1561 cases/100 000 in 2012. Monaghan et al. [206] studied the seasonal abundance of *Ae. aegypti* in the United States as a means of estimating Zika virus infection risk [206]. All 50 included cities were found to be suitable during the summer months (July to September), while only cities in Florida and Texas were found to have *Ae. aegypti* abundance potential during the winter months (December to March). Lo and Park [207] found that regions of Brazil with elevated temperature and precipitation were more conducive to *Ae. aegypti* presence and Zika virus cases. Da Cruz Ferreira et al. [20] found that dengue occurrence increased by 25% when the average number of mosquitoes caught by traps increased by 0.1 per week. Stewart-Ibarra and Lowe [208] assessed the effect of climatic and entomological variables on intra-annual variability in dengue incidence in Southern Ecuador. Da Rocha Taranto et al. [209] examined the relationship between vector collection, species composition, hatching rates, and population density on dengue incidence. Hatching rate was found to be affected by population density and climate, and presence of vectors was associated with dengue incidence [209]. Ernst et al. [192] found no correlation between *Ae. aegypti* density and human age structure between two cities with different dengue transmission dynamics.

## Discussion

Several main conclusions can be drawn from the existing literature. Firstly, temperature and humidity have a significant impact on mosquito distribution and dynamics [18–20,22,24], development of immatures [42,43,43,57,58,60], and arboviral infection rates. Minimum temperature in particular appears to have a large impact on mosquito dynamics, including oviposition [36,42] and survival [43]. Photoperiod is also emerging as a determinant of oviposition [46] and survival [23], but more research on this topic is needed.

Low socioeconomic status and related factors, including poor infrastructure, inconsistent access to water, and high household resident density, have been consistently with arbovirus vector occurrence [73–76,90], suggesting that socioeconomic development may be an important means of vector-borne disease prevention.

Effects of interspecific competition on arboviral vector species is currently poorly understood. Among several laboratory experimental studies [31,63–67] and one observational field study [14], a clear picture of the effects of one mosquito species on the population dynamics and ecology of other mosquito species is still absent. More research on this topic is needed. In terms of population dynamics in general, studies indicate that under conditions of climate change, arboviral vector populations will shift towards the poles, resulting in a reduction of vector burden in regions such as Northern Brazil [102], but an increase in vector population density and corresponding vector-borne disease risk in regions such as Argentina [105] and North America [104,106–108,110,111].

To determine vector competence, a species must be able to acquire, maintain, and transmit a pathogen, which is assessed through experimental infection studies. However, these studies are heterogeneous in both the mosquito populations and virus strains used, as well as methods measuring potential to transmit [210] The detection of viral particles in a wild-caught mosquitoes does not signify vector competence on its own, but it lends support to evidence from laboratory studies, when coupled with the observation of human host-feeding behaviour. Field studies are also important to assess the relative importance of competent vector species in disease maintenance and/or transmission. Vector competence for Zika virus has been well established for *Ae. aegypti* [117–130] and *Ae. albopictus* [120,122], but there is a growing consensus that *Cx. quinquefasciatus* is not a competent Zika virus vector, and no consensus has been reached regarding the competence of *Ae vexans*. A number of studies report that *Cx. quinquefasciatus* is refractory to Zika virus [128,134–137,139,211]. While Zika virus has been detected in a small number of field-caught *Cx. quinquefasciatus* in Brazil [140], this does not necessarily indicate their ability to transmit the virus. Interestingly, information on Zika virus vector species composition was abundant, but sparse on Zika virus vector transmission dynamics. Few studies examined natural infection rates of Zika virus [129,130], vertical transmission [179], or co-infection with other viruses [126]. This is in contrast to the wealth of research available on natural infection and co-infection for dengue and chikungunya viruses, although vertical transmission research was sparse for all three viruses [144,148,156,175,178,197,198]. Based on the internationally recognized urgency of Zika virus infection as a public health concern, and potential increase in the importance of this and other emerging arboviruses in the future, further research on arbovirus vectors and transmission dynamics is of pressing need.

## Acknowledgements

This research is part of an international project entitled ‘Research on Arbovirus Dynamics and Mitigation – Latin America and Canada’ (RADAM-LAC), with field study sites in Colombia, Ecuador and Argentina. The RADAM-LAC Research Team consists of Beate Sander, Camila Gonzalez, Manisha A. Kulkarni, Marcos Miretti, Mauricio Espinel, Jianhong Wu and Varsovia Cevallos. This study was funded by a grant from the Canadian Institutes for Health Research (CIHR) and International Development Research Centre (IDRC)’s CIHR-IDRC Canada-Latin America and Caribbean Zika Virus Research Program to the RADAM-LAC Research Team, and an Early Researcher Award from the Ontario Ministry of Research, Innovation and Science to MK.

## Supporting Information Captions

**Table S1. List of full-text articles included in the review**. Information on first author’s last name, year of publication, title, journal, review section, study design, and arbovirus and mosquito vector species of interest are given for each full-text article.

**Table S2. PRISMA-ScR Checklist**. Checklist stating location of each element of the scoping review, as implemented by Tricco et al. [212].

## References

1. Dick GWA, Kitchen SF, Haddow AJ. Zika virus. I. Isolations and serological specificity. Trans R Soc Trop Med Hyg. 1952;46: 509–520.

2. Weaver SC, Costa F, Garcia-Blanco MA, Ko AI, Ribeiro GS, Saade G, et al. Zika virus: History, emergence, biology, and prospects for control. Antiviral Res. 2016;130: 69–80. doi:10.1016/j.antiviral.2016.03.010

3. World Health Organization. Zika virus [Internet]. 20 Jul 2018 [cited 6 Jun 2019]. Available: https://www.who.int/news-room/fact-sheets/detail/zika-virus

4. Hills SL, Fischer M, Petersen LR. Epidemiology of Zika Virus Infection. J Infect Dis. 2017;216: S868–S874. doi:10.1093/infdis/jix434

5. Kamgang B, Yougang AP, Tchoupo M, Riveron JM, Wondji C. Temporal distribution and insecticide resistance profile of two major arbovirus vectors Aedes aegypti and Aedes albopictus in Yaoundé, the capital city of Cameroon. Parasit Vectors. 2017;10: 469. doi:10.1186/s13071-017-2408-x

6. Murray NEA, Quam MB, Wilder-Smith A. Epidemiology of dengue: past, present and future prospects. Clin Epidemiol. 2013;5: 299–309. doi:10.2147/CLEP.S34440

7. Zeller H, Van Bortel W, Sudre B. Chikungunya: Its History in Africa and Asia and Its Spread to New Regions in 2013-2014. J Infect Dis. 2016;214: S436–S440. doi:10.1093/infdis/jiw391

8. Yactayo S, Staples JE, Millot V, Cibrelus L, Ramon-Pardo P. Epidemiology of Chikungunya in the Americas. J Infect Dis. 2016;214: S441–S445. doi:10.1093/infdis/jiw390

9. Centre for Disease Control. Clinical Guidance | Dengue | CDC [Internet]. 15 Jan 2019 [cited 2 May 2019]. Available: https://www.cdc.gov/dengue/clinicallab/clinical.html

10. Nakkhara P, Chongsuvivatwong V, Thammapalo S. Risk factors for symptomatic and asymptomatic chikungunya infection. Trans R Soc Trop Med Hyg. 2013;107: 789–796. doi:10.1093/trstmh/trt083

11. Haby MM, Pinart M, Elias V, Reveiz L. Prevalence of asymptomatic Zika virus infection: a systematic review. Bull World Health Organ. 2018;96: 402–413D. doi:10.2471/BLT.17.201541

12. Krauer F, Riesen M, Reveiz L, Oladapo OT, Martínez-Vega R, Porgo TV, et al. Zika Virus Infection as a Cause of Congenital Brain Abnormalities and Guillain-Barré Syndrome: Systematic Review. PLoS Med. 2017;14: e1002203. doi:10.1371/journal.pmed.1002203

13. Powell JR, Tabachnick WJ, Powell JR, Tabachnick WJ. History of domestication and spread of Aedes aegypti - A Review. Mem Inst Oswaldo Cruz. 2013;108: 11–17. doi:10.1590/0074-0276130395

14. Wagman J, Grieco JP, King R, Briceño I, Bautista K, Polanco J, et al. First record and demonstration of a southward expansion of Aedes albopictus into Orange Walk Town, Belize, Central America. J Am Mosq Control Assoc. 2013;29: 380–382. doi:10.2987/13-6364.1

15. Miller MJ, Loaiza JR. Geographic expansion of the invasive mosquito Aedes albopictus across Panama--implications for control of dengue and Chikungunya viruses. PLoS Negl Trop Dis. 2015;9: e0003383. doi:10.1371/journal.pntd.0003383

16. Waddell LA, Greig JD. Scoping Review of the Zika Virus Literature. PloS One. 2016;11: e0156376. doi:10.1371/journal.pone.0156376

17. Reiskind MH, Lounibos LP. Spatial and temporal patterns of abundance of Aedes aegypti L. (Stegomyia aegypti) and Aedes albopictus (Skuse) [Stegomyia albopictus (Skuse)] in southern Florida. Med Vet Entomol. 2013;27: 421–429. doi:10.1111/mve.12000

18. Degener CM, Ázara TMF de, Roque RA, Codeço CT, Nobre AA, Ohly JJ, et al. Temporal abundance of Aedes aegypti in Manaus, Brazil, measured by two trap types for adult mosquitoes. Mem Inst Oswaldo Cruz. 2014;109: 1030–1040. doi:10.1590/0074-0276140234

19. Lucio PS, Degallier N, Servain J, Hannart A, Durand B, de Souza RN, et al. A case study of the influence of local weather on Aedes aegypti (L.) aging and mortality. J Vector Ecol J Soc Vector Ecol. 2013;38: 20–37. doi:10.1111/j.1948-7134.2013.12005.x

20. da Cruz Ferreira DA, Degener CM, de Almeida Marques-Toledo C, Bendati MM, Fetzer LO, Teixeira CP, et al. Meteorological variables and mosquito monitoring are good predictors for infestation trends of Aedes aegypti, the vector of dengue, chikungunya and Zika. Parasit Vectors. 2017;10: 78. doi:10.1186/s13071-017-2025-8

21. Regis LN, Acioli RV, Silveira JC, de Melo-Santos MAV, da Cunha MCS, Souza F, et al. Characterization of the spatial and temporal dynamics of the dengue vector population established in urban areas of Fernando de Noronha, a Brazilian oceanic island. Acta Trop. 2014;137: 80–87. doi:10.1016/j.actatropica.2014.04.010

22. Goindin D, Delannay C, Ramdini C, Gustave J, Fouque F. Parity and longevity of Aedes aegypti according to temperatures in controlled conditions and consequences on dengue transmission risks. PloS One. 2015;10: e0135489. doi:10.1371/journal.pone.0135489

23. Costanzo KS, Schelble S, Jerz K, Keenan M. The effect of photoperiod on life history and blood-feeding activity in Aedes albopictus and Aedes aegypti (Diptera: Culicidae). J Vector Ecol J Soc Vector Ecol. 2015;40: 164–171. doi:10.1111/jvec.12146

24. Lega J, Brown HE, Barrera R. Aedes aegypti (Diptera: Culicidae) Abundance Model Improved With Relative Humidity and Precipitation-Driven Egg Hatching. J Med Entomol. 2017;54: 1375–1384. doi:10.1093/jme/tjx077

25. Padilla-Torres SD, Ferraz G, Luz SLB, Zamora-Perea E, Abad-Franch F. Modeling dengue vector dynamics under imperfect detection: three years of site-occupancy by Aedes aegypti and Aedes albopictus in urban Amazonia. PloS One. 2013;8: e58420. doi:10.1371/journal.pone.0058420

26. Sanavria A, Silva CB da, Electo ãH, Nogueira LCR, Thomé SMG, Angelo I da C, et al. Intelligent monitoring of Aedes aegypti in a rural area of Rio de Janeiro State, Brazil. Rev Inst Med Trop Sao Paulo. 2017;59: e51. doi:10.1590/S1678-9946201759051

27. Simões TC, Codeço CT, Nobre AA, Eiras AE. Modeling the non-stationary climate dependent temporal dynamics of Aedes aegypti. PloS One. 2013;8: e64773. doi:10.1371/journal.pone.0064773

28. Faraji A, Egizi A, Fonseca DM, Unlu I, Crepeau T, Healy SP, et al. Comparative host feeding patterns of the Asian tiger mosquito, Aedes albopictus, in urban and suburban Northeastern USA and implications for disease transmission. PLoS Negl Trop Dis. 2014;8: e3037. doi:10.1371/journal.pntd.0003037

29. Vinauger C, Lahondère C, Wolff GH, Locke LT, Liaw JE, Parrish JZ, et al. Modulation of Host Learning in Aedes aegypti Mosquitoes. Curr Biol CB. 2018;28: 333–344.e8. doi:10.1016/j.cub.2017.12.015

30. Fader JE, Juliano SA. Oviposition habitat selection by container-dwelling mosquitoes: responses to cues of larval and detritus abundances in the field. Ecol Entomol. 2014;39: 245–252. doi:10.1111/een.12095

31. Allgood DW, Yee DA. Oviposition preference and offspring performance in container breeding mosquitoes: evaluating the effects of organic compounds and laboratory colonisation. Ecol Entomol. 2017;42: 506–516. doi:10.1111/een.12412

32. Fonseca DM, Kaplan LR, Heiry RA, Strickman D. Density-Dependent Oviposition by Female Aedes albopictus (Diptera: Culicidae) Spreads Eggs Among Containers During the Summer but Accumulates Them in the Fall. J Med Entomol. 2015;52: 705–712. doi:10.1093/jme/tjv060

33. Chitolina RF, Anjos FA, Lima TS, Castro EA, Costa-Ribeiro MCV. Raw sewage as breeding site to Aedes (Stegomyia) aegypti (Diptera, culicidae). Acta Trop. 2016;164: 290–296. doi:10.1016/j.actatropica.2016.07.013

34. Williges E, Faraji A, Gaugler R. Vertical Oviposition Preferences of the Asian Tiger Mosquito, Aedes albopictus, In Temperate North America. J Am Mosq Control Assoc. 2014;30: 169–174. doi:10.2987/14-6409R.1

35. Rey JR, O’Connell SM. Oviposition by Aedes aegypti and Aedes albopictus: influence of congeners and of oviposition site characteristics. J Vector Ecol J Soc Vector Ecol. 2014;39: 190–196. doi:10.1111/j.1948-7134.2014.12086.x

36. Brown HE, Cox J, Comrie AC, Barrera R. Habitat and Density of Oviposition Opportunity Influences Aedes aegypti (Diptera: Culicidae) Flight Distance. J Med Entomol. 2017;54: 1385–1389. doi:10.1093/jme/tjx083

37. Santos ND de L, Paixão K da S, Napoleão TH, Trindade PB, Pinto MR, Coelho LCBB, et al. Evaluation of Moringa oleifera seed lectin in traps for the capture of Aedes aegypti eggs and adults under semi-field conditions. Parasitol Res. 2014;113: 1837–1842. doi:10.1007/s00436-014-3830-z

38. Wermelinger ED, Ferreira AP, Carvalho RW de, Silva AA da, Benigno CV. Aedes aegypti eggs oviposited on water surface collected from field ovitraps in Nova Iguaçu City, Brazil. Rev Soc Bras Med Trop. 2015;48: 770–772. doi:10.1590/0037-8682-0087-2015

39. Valença MA, Marteis LS, Steffler LM, Silva AM, Santos RLC. Dynamics and characterization of Aedes aegypti (L.) (Diptera: Culicidae) key breeding sites. Neotrop Entomol. 2013;42: 311–316. doi:10.1007/s13744-013-0118-4

40. Chadee DD, Martinez R. Aedes aegypti (L.) in Latin American and Caribbean region: With growing evidence for vector adaptation to climate change? Acta Trop. 2016;156: 137–143. doi:10.1016/j.actatropica.2015.12.022

41. Cavalcanti LP de G, Oliveira R de MAB, Alencar CH. Changes in infestation sites of female Aedes aegypti in Northeast Brazil. Rev Soc Bras Med Trop. 2016;49: 498–501. doi:10.1590/0037-8682-0044-2016

42. Estallo EL, Ludueña-Almeida FF, Introini MV, Zaidenberg M, Almirón WR. Weather Variability Associated with Aedes (Stegomyia) aegypti (Dengue Vector) Oviposition Dynamics in Northwestern Argentina. PloS One. 2015;10: e0127820. doi:10.1371/journal.pone.0127820

43. De Majo MS, Montini P, Fischer S. Egg Hatching and Survival of Immature Stages of Aedes aegypti (Diptera: Culicidae) Under Natural Temperature Conditions During the Cold Season in Buenos Aires, Argentina. J Med Entomol. 2017;54: 106–113. doi:10.1093/jme/tjw131

44. Resende MC de, Silva IM, Ellis BR, Eiras ÁE. A comparison of larval, ovitrap and MosquiTRAP surveillance for Aedes (Stegomyia) aegypti. Mem Inst Oswaldo Cruz. 2013;108: 1024–1030. doi:10.1590/0074-0276130128

45. Dibo MR, Fávaro EA, Parra MCP, Santos TC dos, Cassiano JH, Deitz KV de S, et al. Evaluation of two sweeping methods for estimating the number of immature Aedes aegypti (Diptera: Culicidae) in large containers. Rev Soc Bras Med Trop. 2013;46: 502–505. doi:10.1590/0037-8682-1432-2013

46. Farnesi LC, Barbosa CS, Araripe LO, Bruno RV. The influence of a light and dark cycle on the egg laying activity of Aedes aegypti (Linnaeus, 1762) (Diptera: Culicidae). Mem Inst Oswaldo Cruz. 2018;113: e170362. doi:10.1590/0074-02760170362

47. Albeny-Simões D, Murrell EG, Elliot SL, Andrade MR, Lima E, Juliano SA, et al. Attracted to the enemy: Aedes aegypti prefers oviposition sites with predator-killed conspecifics. Oecologia. 2014;175: 481–492. doi:10.1007/s00442-014-2910-1

48. Wasserberg G, White L, Bullard A, King J, Maxwell R. Oviposition site selection in Aedes albopictus (Diptera: Culicidae): are the effects of predation risk and food level independent? J Med Entomol. 2013;50: 1159–1164.

49. Davis TJ, Kaufman PE, Hogsette JA, Kline DL. The Effects of Larval Habitat Quality on Aedes albopictus Skip Oviposition. J Am Mosq Control Assoc. 2015;31: 321–328. doi:10.2987/moco-31-04-321-328.1

50. Abreu FVS de, Morais MM, Ribeiro SP, Eiras ÁE. Influence of breeding site availability on the oviposition behaviour of Aedes aegypti. Mem Inst Oswaldo Cruz. 2015;110: 669–676. doi:10.1590/0074-02760140490

51. Wasserberg G, Bailes N, Davis C, Yeoman K. Hump-shaped density-dependent regulation of mosquito oviposition site-selection by conspecific immature stages: theory, field test with Aedes albopictus, and a meta-analysis. PloS One. 2014;9: e92658. doi:10.1371/journal.pone.0092658

52. Ruktanonchai NW, Lounibos LP, Smith DL, Allan SA. Experience- and age-mediated oviposition behaviour in the yellow fever mosquito Stegomyia aegypti (=Aedes aegypti). Med Vet Entomol. 2015;29: 255–262. doi:10.1111/mve.12119

53. Giménez JO, Fischer S, Zalazar L, Stein M. Cold Season Mortality Under Natural Conditions and Subsequent Hatching Response of Aedes (Stegomyia) aegypti (Diptera: Culicidae) Eggs in a Subtropical City of Argentina. J Med Entomol. 2015;52: 879–885. doi:10.1093/jme/tjv107

54. Byttebier B, De Majo MS, De Majo MS, Fischer S. Hatching response of Aedes aegypti (Diptera: Culicidae) eggs at low temperatures: effects of hatching media and storage conditions. J Med Entomol. 2014;51: 97–103.

55. Grech MG, Sartor PD, Almirón WR, Ludueña-Almeida FF. Effect of temperature on life history traits during immature development of Aedes aegypti and Culex quinquefasciatus (Diptera: Culicidae) from Córdoba city, Argentina. Acta Trop. 2015;146: 1–6. doi:10.1016/j.actatropica.2015.02.010

56. Marinho RA, Beserra EB, Bezerra-Gusmão MA, Porto V de S, Olinda RA, Dos Santos CAC. Effects of temperature on the life cycle, expansion, and dispersion of Aedes aegypti (Diptera: Culicidae) in three cities in Paraiba, Brazil. J Vector Ecol J Soc Vector Ecol. 2016;41: 1–10. doi:10.1111/jvec.12187

57. De Majo MS, Fischer S, Otero M, Schweigmann N. Effects of thermal heterogeneity and egg mortality on differences in the population dynamics of Aedes aegypti (Diptera: Culicidae) over short distances in temperate Argentina. J Med Entomol. 2013;50: 543–551.

58. Couret J, Dotson E, Benedict MQ. Temperature, larval diet, and density effects on development rate and survival of Aedes aegypti (Diptera: Culicidae). PloS One. 2014;9: e87468. doi:10.1371/journal.pone.0087468

59. Lang BJ, Idugboe S, McManus K, Drury F, Qureshi A, Cator LJ. The Effect of Larval Diet on Adult Survival, Swarming Activity and Copulation Success in Male Aedes aegypti (Diptera: Culicidae). J Med Entomol. 2018;55: 29–35. doi:10.1093/jme/tjx187

60. Lopes TF, Holcman MM, Barbosa GL, Domingos M de F, Barreiros RMOV. Laboratory evaluation of the development of Aedes aegypti in two seasons: influence of different places and different densities. Rev Inst Med Trop Sao Paulo. 2014;56: 369–374.

61. de Brito Arduino M, Mucci LF, Serpa LLN, Rodrigues M de M. Effect of salinity on the behavior of Aedes aegypti populations from the coast and plateau of southeastern Brazil. J Vector Borne Dis. 2015;52: 79–87.

62. Romeo Aznar V, De Majo MS, Fischer S, Francisco D, Natiello MA, Solari HG. A model for the development of Aedes (Stegomyia) aegypti as a function of the available food. J Theor Biol. 2015;365: 311–324. doi:10.1016/j.jtbi.2014.10.016

63. Noden BH, O’Neal PA, Fader JE, Juliano SA. Impact of inter- and intra-specific competition among larvae on larval, adult, and life-table traits ofAedes aegyptiandAedes albopictusfemales. Ecol Entomol. 2016;41: 192–200. doi:10.1111/een.12290

64. Yee DA, Skiff JF. Interspecific competition of a new invasive mosquito, Culex coronator, and two container mosquitoes, Aedes albopictus and Cx. quinquefasciatus (Diptera: Culicidae), across different detritus environments. J Med Entomol. 2014;51: 89–96.

65. Riback TIS, Honório NA, Pereira RN, Godoy WAC, Codeço CT. Better to Be in Bad Company than to Be Alone? Aedes Vectors Respond Differently to Breeding Site Quality in the Presence of Others. PloS One. 2015;10: e0134450. doi:10.1371/journal.pone.0134450

66. Alto BW, Bettinardi DJ, Ortiz S. Interspecific Larval Competition Differentially Impacts Adult Survival in Dengue Vectors. J Med Entomol. 2015;52: 163–170. doi:10.1093/jme/tju062

67. Kesavaraju B, Leisnham PT, Keane S, Delisi N, Pozatti R. Interspecific Competition between Aedes albopictus and A. sierrensis: potential for Competitive Displacement in the Western United States. PloS One. 2014;9: e89698. doi:10.1371/journal.pone.0089698

68. Jaimes-Dueñez J, Arboleda S, Triana-Chávez O, Gómez-Palacio A. Spatio-temporal distribution of Aedes aegypti (Diptera: Culicidae) mitochondrial lineages in cities with distinct dengue incidence rates suggests complex population dynamics of the dengue vector in Colombia. PLoS Negl Trop Dis. 2015;9: e0003553. doi:10.1371/journal.pntd.0003553

69. Steffler LM, Dolabella SS, Ribolla PEM, Dreyer CS, Araújo ED, Oliveira RG, et al. Genetic variability and spatial distribution in small geographic scale of Aedes aegypti (Diptera: Culicidae) under different climatic conditions in Northeastern Brazil. Parasit Vectors. 2016;9: 530. doi:10.1186/s13071-016-1814-9

70. Kotsakiozi P, Gloria-Soria A, Caccone A, Evans B, Schama R, Martins AJ, et al. Tracking the return of Aedes aegypti to Brazil, the major vector of the dengue, chikungunya and Zika viruses. PLoS Negl Trop Dis. 2017;11: e0005653. doi:10.1371/journal.pntd.0005653

71. Guagliardo SA, Morrison AC, Luis Barboza J, Wesson DM, Ponnusamy L, Astete H, et al. Evidence for Aedes aegypti (Diptera: Culicidae) Oviposition on Boats in the Peruvian Amazon. J Med Entomol. 2015;52: 726–729. doi:10.1093/jme/tjv048

72. Guagliardo SA, Morrison AC, Barboza JL, Requena E, Astete H, Vazquez-Prokopec G, et al. River boats contribute to the regional spread of the dengue vector Aedes aegypti in the Peruvian Amazon. PLoS Negl Trop Dis. 2015;9: e0003648. doi:10.1371/journal.pntd.0003648

73. Rodrigues M de M, Marques GRAM, Serpa LLN, Arduino M de B, Voltolini JC, Barbosa GL, et al. Density of Aedes aegypti and Aedes albopictus and its association with number of residents and meteorological variables in the home environment of dengue endemic area, São Paulo, Brazil. Parasit Vectors. 2015;8: 115. doi:10.1186/s13071-015-0703-y

74. Favaro EA, Dibo MR, Pereira M, Chierotti AP, Rodrigues-Junior AL, Chiaravalloti-Neto F. Aedes aegypti entomological indices in an endemic area for dengue in Sao Paulo State, Brazil. Rev Saude Publica. 2013;47: 588–597.

75. Morales-Pérez A, Nava-Aguilera E, Balanzar-Martínez A, Cortés-Guzmán AJ, Gasga-Salinas D, Rodríguez-Ramos IE, et al. Aedes aegypti breeding ecology in Guerrero: crosssectional study of mosquito breeding sites from the baseline for the Camino Verde trial in Mexico. BMC Public Health. 2017;17: 450. doi:10.1186/s12889-017-4293-9

76. Schafrick NH, Milbrath MO, Berrocal VJ, Wilson ML, Eisenberg JNS. Spatial clustering of Aedes aegypti related to breeding container characteristics in Coastal Ecuador: implications for dengue control. Am J Trop Med Hyg. 2013;89: 758–765. doi:10.4269/ajtmh.12-0485

77. Serpa LLN, Monteiro Marques GRA, de Lima AP, Voltolini JC, Arduino M de B, Barbosa GL, et al. Study of the distribution and abundance of the eggs of Aedes aegypti and Aedes albopictus according to the habitat and meteorological variables, municipality of São Sebastião, São Paulo State, Brazil. Parasit Vectors. 2013;6: 321. doi:10.1186/1756-3305-6-321

78. Arana-Guardia R, Baak-Baak CM, Loroño-Pino MA, Machain-Williams C, Beaty BJ, Eisen L, et al. Stormwater drains and catch basins as sources for production of Aedes aegypti and Culex quinquefasciatus. Acta Trop. 2014;134: 33–42. doi:10.1016/j.actatropica.2014.01.011

79. Paploski IAD, Rodrigues MS, Mugabe VA, Kikuti M, Tavares AS, Reis MG, et al. Storm drains as larval development and adult resting sites for Aedes aegypti and Aedes albopictus in Salvador, Brazil. Parasit Vectors. 2016;9: 419. doi:10.1186/s13071-016-1705-0

80. Baak-Baak CM, Arana-Guardia R, Cigarroa-Toledo N, Puc-Tinal M, Coba-Tún C, Rivero-Osorno V, et al. Urban Mosquito Fauna in Mérida City, México: Immatures Collected from Containers and Storm-water Drains/Catch Basins. Southwest Entomol. 2014;39: 291–306. doi:10.3958/059.039.0207

81. Overgaard HJ, Olano VA, Jaramillo JF, Matiz MI, Sarmiento D, Stenström TA, et al. A cross-sectional survey of Aedes aegypti immature abundance in urban and rural household containers in central Colombia. Parasit Vectors. 2017;10: 356. doi:10.1186/s13071-017-2295-1

82. Unlu I, Faraji A, Indelicato N, Fonseca DM. The hidden world of Asian tiger mosquitoes: immature Aedes albopictus (Skuse) dominate in rainwater corrugated extension spouts. Trans R Soc Trop Med Hyg. 2014;108: 699–705. doi:10.1093/trstmh/tru139

83. Samson DM, Qualls WA, Roque D, Naranjo DP, Alimi T, Arheart KL, et al. Resting and energy reserves of Aedes albopictus collected in common landscaping vegetation in St. Augustine, Florida. J Am Mosq Control Assoc. 2013;29: 231–236. doi:10.2987/13-6347R.1

84. Quintero J, Brochero H, Manrique-Saide P, Barrera-Pérez M, Basso C, Romero S, et al. Ecological, biological and social dimensions of dengue vector breeding in five urban settings of Latin America: a multi-country study. BMC Infect Dis. 2014;14: 38. doi:10.1186/1471-2334-14-38

85. Yee DA, Abuzeineh AA, Ezeakacha NF, Schelble SS, Glasgow WC, Flanagan SD, et al. Mosquito Larvae in Tires from Mississippi, United States: The Efficacy of Abiotic and Biotic Parameters in Predicting Spatial and Temporal Patterns of Mosquito Populations and Communities. J Med Entomol. 2015;52: 394–407. doi:10.1093/jme/tjv028

86. Wilke ABB, Medeiros-Sousa AR, Ceretti-Junior W, Marrelli MT. Mosquito populations dynamics associated with climate variations. Acta Trop. 2017;166: 343–350. doi:10.1016/j.actatropica.2016.10.025

87. Montagner FRG, Silva OS, Jahnke SM. Mosquito species occurrence in association with landscape composition in green urban areas. Braz J Biol Rev Brasleira Biol. 2017; 0. doi:10.1590/1519-6984.04416

88. Hagenlocher M, Delmelle E, Casas I, Kienberger S. Assessing socioeconomic vulnerability to dengue fever in Cali, Colombia: statistical vs expert-based modeling. Int J Health Geogr. 2013;12: 36. doi:10.1186/1476-072X-12-36

89. Silva Lima AW, Honório NA, Codeço CT. Spatial Clustering of Aedes aegypti (Diptera: Culicidae) and Its Impact on Entomological Surveillance Indicators. J Med Entomol. 2016;53: 343–348. doi:10.1093/jme/tjv187

90. Fuentes-Vallejo M, Higuera-Mendieta DR, García-Betancourt T, Alcalá-Espinosa LA, García-Sánchez D, Munévar-Cagigas DA, et al. Territorial analysis of Aedes aegypti distribution in two Colombian cities: a chorematic and ecosystem approach. Cad Saude Publica. 2015;31: 517–530.

91. Manrique-Saide P, Davies CR, Coleman PG, Che-Mendoza A, Dzul-Manzanilla F, Barrera-Pérez M, et al. The risk of Aedes aegypti breeding and premises condition in South Mexico. J Am Mosq Control Assoc. 2013;29: 337–345. doi:10.2987/13-6350.1

92. Espinosa MO, Polop F, Rotela CH, Abril M, Scavuzzo CM. Spatial pattern evolution of Aedes aegypti breeding sites in an Argentinean city without a dengue vector control programme. Geospatial Health. 2016;11: 471. doi:10.4081/gh.2016.471

93. Little E, Biehler D, Leisnham PT, Jordan R, Wilson S, LaDeau SL. Socio-Ecological Mechanisms Supporting High Densities of Aedes albopictus (Diptera: Culicidae) in Baltimore, MD. J Med Entomol. 2017;54: 1183–1192. doi:10.1093/jme/tjx103

94. Taber ED, Hutchinson ML, Smithwick EAH, Blanford JI. A decade of colonization: the spread of the Asian tiger mosquito in Pennsylvania and implications for disease risk. J Vector Ecol J Soc Vector Ecol. 2017;42: 3–12. doi:10.1111/jvec.12234

95. Hahn MB, Eisen L, McAllister J, Savage HM, Mutebi J-P, Eisen RJ. Updated Reported Distribution of Aedes (Stegomyia) aegypti and Aedes (Stegomyia) albopictus (Diptera: Culicidae) in the United States, 1995-2016. J Med Entomol. 2017;54: 1420–1424. doi:10.1093/jme/tjx088

96. Hopperstad KA, Reiskind MH. Recent Changes in the Local Distribution of Aedes aegypti (Diptera: Culicidae) in South Florida, USA. J Med Entomol. 2016;53: 836–842. doi:10.1093/jme/tjw050

97. Johnson TL, Haque U, Monaghan AJ, Eisen L, Hahn MB, Hayden MH, et al. Modeling the Environmental Suitability for Aedes (Stegomyia) aegypti and Aedes (Stegomyia) albopictus (Diptera: Culicidae) in the Contiguous United States. J Med Entomol. 2017;54: 1605–1614. doi:10.1093/jme/tjx163

98. Unlu I, Farajollahi A. A multiyear surveillance for Aedes albopictus with Biogents Sentinel trap counts for males and species composition of other mosquito species. J Am Mosq Control Assoc. 2014;30: 122–125. doi:10.2987/14-6401.1

99. Leal-Santos FA, Santana MB de A, de Figueiredo DA, de Oliveira MM, Acel AM, Ribeiro ALM, et al. Effective surveillance of vector dynamics of Aedes aegypti in a hospital setting in Cuiabá, Mato Grosso, Brazil. J Infect Dev Ctries. 2014;8: 1356–1360.

100. Carvalho RG, Lourenço-de-Oliveira R, Braga IA. Updating the geographical distribution and frequency of Aedes albopictus in Brazil with remarks regarding its range in the Americas. Mem Inst Oswaldo Cruz. 2014;109: 787–796.

101. LaCon G, Morrison AC, Astete H, Stoddard ST, Paz-Soldan VA, Elder JP, et al. Shifting patterns of Aedes aegypti fine scale spatial clustering in Iquitos, Peru. PLoS Negl Trop Dis. 2014;8: e3038. doi:10.1371/journal.pntd.0003038

102. Cardoso-Leite R, Vilarinho AC, Novaes MC, Tonetto AF, Vilardi GC, Guillermo-Ferreira R. Recent and future environmental suitability to dengue fever in Brazil using species distribution model. Trans R Soc Trop Med Hyg. 2014;108: 99–104. doi:10.1093/trstmh/trt115

103. Escobar LE, Romero-Alvarez D, Leon R, Lepe-Lopez MA, Craft ME, Borbor-Cordova MJ, et al. Declining Prevalence of Disease Vectors Under Climate Change. Sci Rep. 2016;6: 39150. doi:10.1038/srep39150

104. Estrada-Contreras I, Sandoval-Ruiz CA, Mendoza-Palmero FS, Ibáñez-Bernal S, Equihua M, Benítez G. Data documenting the potential distribution of Aedes aegypti in the center of Veracruz, Mexico. Data Brief. 2017;10: 432–437. doi:10.1016/j.dib.2016.12.014

105. Fischer S, De Majo MS, Quiroga L, Paez M, Schweigmann N. Long-term spatio-temporal dynamics of the mosquito Aedes aegypti in temperate Argentina. Bull Entomol Res. 2017;107: 225–233. doi:10.1017/S0007485316000869

106. Ogden NH, Milka R, Caminade C, Gachon P. Recent and projected future climatic suitability of North America for the Asian tiger mosquito Aedes albopictus. Parasit Vectors. 2014;7: 532. doi:10.1186/s13071-014-0532-4

107. Liu-Helmersson J, Stenlund H, Wilder-Smith A, Rocklöv J. Vectorial capacity of Aedes aegypti: effects of temperature and implications for global dengue epidemic potential. PloS One. 2014;9: e89783. doi:10.1371/journal.pone.0089783

108. Rochlin I, Ninivaggi DV, Hutchinson ML, Farajollahi A. Climate change and range expansion of the Asian tiger mosquito (Aedes albopictus) in Northeastern USA: implications for public health practitioners. PloS One. 2013;8: e60874. doi:10.1371/journal.pone.0060874

109. Hahn MB, Eisen RJ, Eisen L, Boegler KA, Moore CG, McAllister J, et al. Reported Distribution of Aedes (Stegomyia) aegypti and Aedes (Stegomyia) albopictus in the United States, 1995-2016 (Diptera: Culicidae). J Med Entomol. 2016; doi:10.1093/jme/tjw072

110. Morin CW, Comrie AC, Ernst K. Climate and dengue transmission: evidence and implications. Environ Health Perspect. 2013;121: 1264–1272. doi:10.1289/ehp.1306556

111. Ng V, Fazil A, Gachon P, Deuymes G, Radojević M, Mascarenhas M, et al. Assessment of the Probability of Autochthonous Transmission of Chikungunya Virus in Canada under Recent and Projected Climate Change. Environ Health Perspect. 2017;125: 067001. doi:10.1289/EHP669

112. Samson DM, Archer RS, Alimi TO, Arheart KL, Impoinvil DE, Oscar R, et al. New baseline environmental assessment of mosquito ecology in northern Haiti during increased urbanization. J Vector Ecol J Soc Vector Ecol. 2015;40: 46–58. doi:10.1111/jvec.12131

113. Ceretti-Junior W, de Oliveira Christe R, Rizzo M, Strobel RC, de Matos Junior MO, de Mello MHSH, et al. Species Composition and Ecological Aspects of Immature Mosquitoes (Diptera: Culicidae) in Bromeliads in Urban Parks in the City of São Paulo, Brazil. J Arthropod-Borne Dis. 2016;10: 102–112.

114. Ceretti-Júnior W, Medeiros-Sousa AR, Multini LC, Urbinatti PR, Vendrami DP, Natal D, et al. Immature mosquitoes in bamboo internodes in municipal parks, city of são paulo, Brazil. J Am Mosq Control Assoc. 2014;30: 268–274. doi:10.2987/14-6403R.1

115. Lima-Camara TN, Urbinatti PR, Chiaravalloti-Neto F. Finding Aedes aegypti in a natural breeding site in an urban zone, Sao Paulo, Southeastern Brazil. Rev Saude Publica. 2016;50: 3. doi:10.1590/S1518-8787.2016050006245

116. Zanotti G, De Majo MS, Alem I, Schweigmann N, Campos RE, Fischer S. New records of Aedes aegypti at the southern limit of its distribution in Buenos Aires province, Argentina. J Vector Ecol J Soc Vector Ecol. 2015;40: 408–411. doi:10.1111/jvec.12181

117. Bolling BG, Vasilakis N, Guzman H, Widen SG, Wood TG, Popov VL, et al. Insect-specific viruses detected in laboratory mosquito colonies and their potential implications for experiments evaluating arbovirus vector competence. Am J Trop Med Hyg. 2015;92: 422– 428. doi:10.4269/ajtmh.14-0330

118. Azar SR, Roundy CM, Rossi SL, Huang JH, Leal G, Yun R, et al. Differential Vector Competency ofAedes albopictusPopulations from the Americas for Zika Virus. Am J Trop Med Hyg. 2017;97: 330–339. doi:10.4269/ajtmh.16-0969

119. Costa-da-Silva AL, Ioshino RS, Araújo HRC de, Kojin BB, Zanotto PM de A, Oliveira DBL, et al. Correction: Laboratory strains of Aedes aegypti are competent to Brazilian Zika virus. PloS One. 2017;12: e0174081. doi:10.1371/journal.pone.0174081

120. Chouin-Carneiro T, Vega-Rua A, Vazeille M, Yebakima A, Girod R, Goindin D, et al. Differential Susceptibilities of Aedes aegypti and Aedes albopictus from the Americas to Zika Virus. PLoS Negl Trop Dis. 2016;10: e0004543. doi:10.1371/journal.pntd.0004543

121. Cevallos V, Ponce P, Waggoner JJ, Pinsky BA, Coloma J, Quiroga C, et al. Zika and Chikungunya virus detection in naturally infected Aedes aegypti in Ecuador. Acta Trop. 2018;177: 74–80. doi:10.1016/j.actatropica.2017.09.029

122. Ciota AT, Bialosuknia SM, Zink SD, Brecher M, Ehrbar DJ, Morrissette MN, et al. Effects of Zika Virus Strain and Aedes Mosquito Species on Vector Competence. Emerg Infect Dis. 2017;23: 1110–1117. doi:10.3201/eid2307.161633

123. Huang Y-JS, Lyons AC, Hsu W-W, Park SL, Higgs S, Vanlandingham DL. Differential outcomes of Zika virus infection in Aedes aegypti orally challenged with infectious blood meals and infectious protein meals. PloS One. 2017;12: e0182386. doi:10.1371/journal.pone.0182386

124. O’Donnell KL, Bixby MA, Morin KJ, Bradley DS, Vaughan JA. Potential of a Northern Population of Aedes vexans (Diptera: Culicidae) to Transmit Zika Virus. J Med Entomol. 2017;54: 1354–1359. doi:10.1093/jme/tjx087

125. Roundy CM, Azar SR, Rossi SL, Huang JH, Leal G, Yun R, et al. Variation in Aedes aegypti Mosquito Competence for Zika Virus Transmission. Emerg Infect Dis. 2017;23: 625– 632. doi:10.3201/eid2304.161484

126. Rückert C, Weger-Lucarelli J, Garcia-Luna SM, Young MC, Byas AD, Murrieta RA, et al. Impact of simultaneous exposure to arboviruses on infection and transmission by Aedes aegypti mosquitoes. Nat Commun. 2017;8: 15412. doi:10.1038/ncomms15412

127. Secundino NFC, Chaves BA, Orfano AS, Silveira KRD, Rodrigues NB, Campolina TB, et al. Zika virus transmission to mouse ear by mosquito bite: a laboratory model that replicates the natural transmission process. Parasit Vectors. 2017;10: 346. doi:10.1186/s13071-017-2286-2

128. Weger-Lucarelli J, Rückert C, Chotiwan N, Nguyen C, Garcia Luna SM, Fauver JR, et al. Vector Competence of American Mosquitoes for Three Strains of Zika Virus. PLoS Negl Trop Dis. 2016;10: e0005101. doi:10.1371/journal.pntd.0005101

129. Ferreira-de-Brito A, Ribeiro IP, Miranda RM de, Fernandes RS, Campos SS, Silva KAB da, et al. First detection of natural infection of Aedes aegypti with Zika virus in Brazil and throughout South America. Mem Inst Oswaldo Cruz. 2016;111: 655–658. doi:10.1590/0074-02760160332

130. Ayllón T, Campos R de M, Brasil P, Morone FC, Câmara DCP, Meira GLS, et al. Early Evidence for Zika Virus Circulation among Aedes aegypti Mosquitoes, Rio de Janeiro, Brazil. Emerg Infect Dis. 2017;23: 1411–1412. doi:10.3201/eid2308.162007

131. Gendernalik A, Weger-Lucarelli J, Garcia Luna SM, Fauver JR, Rückert C, Murrieta RA, et al. AmericanAedes vexansMosquitoes are Competent Vectors of Zika Virus. Am J Trop Med Hyg. 2017;96: 1338–1340. doi:10.4269/ajtmh.16-0963

132. Evans MV, Dallas TA, Han BA, Murdock CC, Drake JM. Data-driven identification of potential Zika virus vectors. eLife. 2017;6. doi:10.7554/eLife.22053

133. Althouse BM, Vasilakis N, Sall AA, Diallo M, Weaver SC, Hanley KA. Potential for Zika Virus to Establish a Sylvatic Transmission Cycle in the Americas. PLoS Negl Trop Dis. 2016;10: e0005055. doi:10.1371/journal.pntd.0005055

134. Fernandes RS, Campos SS, Ribeiro PS, Raphael LM, Bonaldo MC, Lourenço-de- Oliveira R. Culex quinquefasciatus from areas with the highest incidence of microcephaly associated with Zika virus infections in the Northeast Region of Brazil are refractory to the virus. Mem Inst Oswaldo Cruz. 2017;112: 577–579. doi:10.1590/0074-02760170145

135. Fernandes RS, Campos SS, Ferreira-de-Brito A, Miranda RM de, Barbosa da Silva KA, Castro MG de, et al. Culex quinquefasciatus from Rio de Janeiro Is Not Competent to Transmit the Local Zika Virus. PLoS Negl Trop Dis. 2016;10: e0004993. doi:10.1371/journal.pntd.0004993

136. Hart CE, Roundy CM, Azar SR, Huang JH, Yun R, Reynolds E, et al. Zika Virus Vector Competency of Mosquitoes, Gulf Coast, United States. Emerg Infect Dis. 2017;23: 559–560. doi:10.3201/eid2303.161636

137. Kenney JL, Romo H, Duggal NK, Tzeng W-P, Burkhalter KL, Brault AC, et al. Transmission Incompetence ofCulex quinquefasciatusandCulex pipiens pipiensfrom North America for Zika Virus. Am J Trop Med Hyg. 2017;96: 1235–1240. doi:10.4269/ajtmh.16-0865

138. Lourenço J, Maia de Lima M, Faria NR, Walker A, Kraemer MU, Villabona-Arenas CJ, et al. Epidemiological and ecological determinants of Zika virus transmission in an urban setting. eLife. 2017;6. doi:10.7554/eLife.29820

139. Dodson BL, Rasgon JL. Vector competence of Anopheles and Culex mosquitoes for Zika virus. PeerJ. 2017;5: e3096. doi:10.7717/peerj.3096

140. Guedes DR, Paiva MH, Donato MM, Barbosa PP, Krokovsky L, Rocha SWDS, et al. Zika virus replication in the mosquito Culex quinquefasciatus in Brazil. Emerg Microbes Infect. 2017;6: e69. doi:10.1038/emi.2017.59

141. Aliota MT, Peinado SA, Osorio JE, Bartholomay LC. Culex pipiens and Aedes triseriatus Mosquito Susceptibility to Zika Virus. Emerg Infect Dis. 2016;22: 1857–1859. doi:10.3201/eid2210.161082

142. Bara J, Rapti Z, Cáceres CE, Muturi EJ. Effect of Larval Competition on Extrinsic Incubation Period and Vectorial Capacity of Aedes albopictus for Dengue Virus. PloS One. 2015;10: e0126703. doi:10.1371/journal.pone.0126703

143. Buckner EA, Alto BW, Lounibos LP. Larval Temperature-Food Effects on Adult Mosquito Infection and Vertical Transmission of Dengue-1 Virus. J Med Entomol. 2016;53: 91–98. doi:10.1093/jme/tjv145

144. Nuckols JT, Huang Y-JS, Higgs S, Miller AL, Pyles RB, Spratt HM, et al. Evaluation of Simultaneous Transmission of Chikungunya Virus and Dengue Virus Type 2 in Infected Aedes aegypti and Aedes albopictus (Diptera: Culicidae). J Med Entomol. 2015;52: 447– 451. doi:10.1093/jme/tjv017

145. Xiao F-Z, Zhang Y, Deng Y-Q, He S, Xie H-G, Zhou X-N, et al. The effect of temperature on the extrinsic incubation period and infection rate of dengue virus serotype 2 infection in Aedes albopictus. Arch Virol. 2014;159: 3053–3057. doi:10.1007/s00705-014-2051-1

146. Lourenço-de-Oliveira R, Rua AV, Vezzani D, Willat G, Vazeille M, Mousson L, et al. Aedes aegypti from temperate regions of South America are highly competent to transmit dengue virus. BMC Infect Dis. 2013;13: 610. doi:10.1186/1471-2334-13-610

147. Hill CL, Sharma A, Shouche Y, Severson DW. Dynamics of midgut microflora and dengue virus impact on life history traits in Aedes aegypti. Acta Trop. 2014;140: 151–157. doi:10.1016/j.actatropica.2014.07.015

148. Muturi EJ, Buckner E, Bara J. Superinfection interference between dengue-2 and dengue-4 viruses in Aedes aegypti mosquitoes. Trop Med Int Health TM IH. 2017;22: 399– 406. doi:10.1111/tmi.12846

149. Dos Santos TP, Cruz OG, da Silva KAB, de Castro MG, de Brito AF, Maspero RC, et al. Dengue serotype circulation in natural populations of Aedes aegypti. Acta Trop. 2017;176: 140–143. doi:10.1016/j.actatropica.2017.07.014

150. Alto BW, Smartt CT, Shin D, Bettinardi D, Malicoate J, Anderson SL, et al. Susceptibility of Florida Aedes aegypti and Aedes albopictus to dengue viruses from Puerto Rico. J Vector Ecol J Soc Vector Ecol. 2014;39: 406–413. doi:10.1111/jvec.12116

151. Gómez-Palacio A, Suaza-Vasco J, Castaño S, Triana O, Uribe S. Aedes albopictus (Skuse, 1894) infected with the American-Asian genotype of dengue type 2 virus in Medellín suggests its possible role as vector of dengue fever in Colombia. Biomed Rev Inst Nac Salud. 2017;37: 135–142.

152. Pérez-Pérez J, Sanabria WH, Restrepo C, Rojo R, Henao E, Triana O, et al. Virological surveillance of Aedes (Stegomyia) aegypti and Aedes (Stegomyia) albopictus as support for decision making for dengue control in Medellín. Biomed Rev Inst Nac Salud. 2017;37: 155–166.

153. Gonçalves CM, Melo FF, Bezerra JMT, Chaves BA, Silva BM, Silva LD, et al. Distinct variation in vector competence among nine field populations of Aedes aegypti from a Brazilian dengue-endemic risk city. Parasit Vectors. 2014;7: 320. doi:10.1186/1756-3305-7-320

154. Serra OP, Cardoso BF, Ribeiro ALM, Santos FAL dos, Slhessarenko RD. Mayaro virus and dengue virus 1 and 4 natural infection in culicids from Cuiabá, state of Mato Grosso, Brazil. Mem Inst Oswaldo Cruz. 2016;111: 20–29. doi:10.1590/0074-02760150270

155. Cruz LC de TA da, Serra OP, Leal-Santos FA, Ribeiro ALM, Slhessarenko RD, Santos MA dos. Natural transovarial transmission of dengue virus 4 in Aedes aegypti from Cuiabá, State of Mato Grosso, Brazil. Rev Soc Bras Med Trop. 2015;48: 18–25. doi:10.1590/0037-8682-0264-2014

156. da Costa CF, Dos Passos RA, Lima JBP, Roque RA, de Souza Sampaio V, Campolina TB, et al. Transovarial transmission of DENV in Aedes aegypti in the Amazon basin: a local model of xenomonitoring. Parasit Vectors. 2017;10: 249. doi:10.1186/s13071-017-2194-5

157. Dzul-Manzanilla F, Martínez NE, Cruz-Nolasco M, Gutiérrez-Castro C, López-Damián L, Ibarra-López J, et al. Arbovirus Surveillance and First Report of Chikungunya Virus in Wild Populations of Aedes aegypti from Guerrero, Mexico. J Am Mosq Control Assoc. 2015;31: 275–277. doi:10.2987/moco-31-03-275-277.1

158. Espinosa M, Giamperetti S, Abril M, Seijo A. Vertical transmission of dengue virus in Aedes aegypti collected in Puerto Iguazú, Misiones, Argentina. Rev Inst Med Trop Sao Paulo. 2014;56: 165–167. doi:10.1590/S0036-46652014000200013

159. Farraudière L, Sonor F, Crico S, ãtienne M, Mousson L, Hamel R, et al. First detection of dengue and chikungunya viruses in natural populations of Aedes aegypti in Martinique during the 2013 - 2015 concomitant outbreak. Rev Panam Salud Publica Pan Am J Public Health. 2017;41: e63.

160. Martínez NE, Dzul-Manzanilla F, Gutiérrez-Castro C, Ibarra-López J, Bibiano-Marín W, López-Damián L, et al. Natural vertical transmission of dengue-1 virus in Aedes aegypti populations in Acapulco, Mexico. J Am Mosq Control Assoc. 2014;30: 143–146. doi:10.2987/14-6402.1

161. Adelman ZN, Anderson MAE, Wiley MR, Murreddu MG, Samuel GH, Morazzani EM, et al. Cooler temperatures destabilize RNA interference and increase susceptibility of disease vector mosquitoes to viral infection. PLoS Negl Trop Dis. 2013;7: e2239. doi:10.1371/journal.pntd.0002239

162. Alto BW, Wiggins K, Eastmond B, Ortiz S, Zirbel K, Lounibos LP. Diurnal Temperature Range and Chikungunya Virus Infection in Invasive Mosquito Vectors. J Med Entomol. 2018;55: 217–224. doi:10.1093/jme/tjx182

163. Dong S, Balaraman V, Kantor AM, Lin J, Grant DG, Held NL, et al. Chikungunya virus dissemination from the midgut of Aedes aegypti is associated with temporal basal lamina degradation during bloodmeal digestion. PLoS Negl Trop Dis. 2017;11: e0005976. doi:10.1371/journal.pntd.0005976

164. Ledermann JP, Borland EM, Powers AM. Minimum infectious dose for chikungunya virus in Aedes aegypti and Ae. albopictus mosquitoes. Rev Panam Salud Publica Pan Am J Public Health. 2017;41: e65.

165. Vega-Rúa A, Lourenço-de-Oliveira R, Mousson L, Vazeille M, Fuchs S, Yébakima A, et al. Chikungunya virus transmission potential by local Aedes mosquitoes in the Americas and Europe. PLoS Negl Trop Dis. 2015;9: e0003780. doi:10.1371/journal.pntd.0003780

166. Vega-Rúa A, Zouache K, Girod R, Failloux A-B, Lourenço-de-Oliveira R. High level of vector competence of Aedes aegypti and Aedes albopictus from ten American countries as a crucial factor in the spread of Chikungunya virus. J Virol. 2014;88: 6294–6306. doi:10.1128/JVI.00370-14

167. Alto BW, Wiggins K, Eastmond B, Velez D, Lounibos LP, Lord CC. Transmission risk of two chikungunya lineages by invasive mosquito vectors from Florida and the Dominican Republic. PLoS Negl Trop Dis. 2017;11: e0005724. doi:10.1371/journal.pntd.0005724

168. Lourenço-de-Oliveira R, Failloux A-B. High risk for chikungunya virus to initiate an enzootic sylvatic cycle in the tropical Americas. PLoS Negl Trop Dis. 2017;11: e0005698. doi:10.1371/journal.pntd.0005698

169. Cigarroa-Toledo N, Blitvich BJ, Cetina-Trejo RC, Talavera-Aguilar LG, Baak-Baak CM, Torres-Chablé OM, et al. Chikungunya Virus in Febrile Humans and Aedes aegypti Mosquitoes, Yucatan, Mexico. Emerg Infect Dis. 2016;22: 1804–1807. doi:10.3201/eid2210.152087

170. Díaz-González EE, Kautz TF, Dorantes-Delgado A, Malo-García IR, Laguna-Aguilar M, Langsjoen RM, et al. First Report of Aedes aegypti Transmission of Chikungunya Virus in the Americas. Am J Trop Med Hyg. 2015;93: 1325–1329. doi:10.4269/ajtmh.15-0450

171. Mordecai EA, Cohen JM, Evans MV, Gudapati P, Johnson LR, Lippi CA, et al. Detecting the impact of temperature on transmission of Zika, dengue, and chikungunya using mechanistic models. PLoS Negl Trop Dis. 2017;11: e0005568. doi:10.1371/journal.pntd.0005568

172. Kang DS, Alcalay Y, Lovin DD, Cunningham JM, Eng MW, Chadee DD, et al. Larval stress alters dengue virus susceptibility in Aedes aegypti (L.) adult females. Acta Trop. 2017;174: 97–101. doi:10.1016/j.actatropica.2017.06.018

173. Maciel-de-Freitas R, Sylvestre G, Gandini M, Koella JC. The influence of dengue virus serotype-2 infection on Aedes aegypti (Diptera: Culicidae) motivation and avidity to blood feed. PloS One. 2013;8: e65252. doi:10.1371/journal.pone.0065252

174. Sylvestre G, Gandini M, Maciel-de-Freitas R. Age-dependent effects of oral infection with dengue virus on Aedes aegypti (Diptera: Culicidae) feeding behavior, survival, oviposition success and fecundity. PloS One. 2013;8: e59933. doi:10.1371/journal.pone.0059933

175. Cecílio SG, Júnior WFS, Tótola AH, de Brito Magalhães CL, Ferreira JMS, de Magalhães JC. Dengue virus detection in Aedes aegypti larvae from southeastern Brazil. J Vector Ecol J Soc Vector Ecol. 2015;40: 71–74. doi:10.1111/jvec.12134

176. Calderón-Arguedas O, Troyo A, Moreira-Soto RD, Marín R, Taylor L. Dengue viruses in Aedes albopictus Skuse from a pineapple plantation in Costa Rica. J Vector Ecol J Soc Vector Ecol. 2015;40: 184–186. doi:10.1111/jvec.12149

177. Pérez-Castro R, Castellanos JE, Olano VA, Matiz MI, Jaramillo JF, Vargas SL, et al. Detection of all four dengue serotypes in Aedes aegypti female mosquitoes collected in a rural area in Colombia. Mem Inst Oswaldo Cruz. 2016;111: 233–240. doi:10.1590/0074-02760150363

178. Buckner EA, Alto BW, Lounibos LP. Vertical transmission of Key West dengue-1 virus by Aedes aegypti and Aedes albopictus (Diptera: Culicidae) mosquitoes from Florida. J Med Entomol. 2013;50: 1291–1297.

179. Thangamani S. Vertical Transmission of Zika Virus in Aedes aegypti Mosquitoes. - PubMed - NCBI. Am J Trop Med Hyg. 2016;95: 1169–1173.

180. Marini G, Guzzetta G, Rosà R, Merler S. First outbreak of Zika virus in the continental United States: a modelling analysis. Euro Surveill Bull Eur Sur Mal Transm Eur Commun Euro0 Euro1. 2017;22. doi:10.2807/1560-7917.ES.2017.22.37.30612

181. Majumder MS, Santillana M, Mekaru SR, McGinnis DP, Khan K, Brownstein JS. Utilizing Nontraditional Data Sources for Near Real-Time Estimation of Transmission Dynamics During the 2015-2016 Colombian Zika Virus Disease Outbreak. JMIR Public Health Surveill. 2016;2: e30. doi:10.2196/publichealth.5814

182. Villela D a. M, Bastos LS, DE Carvalho LM, Cruz OG, Gomes MFC, Durovni B, et al. Zika in Rio de Janeiro: Assessment of basic reproduction number and comparison with dengue outbreaks. Epidemiol Infect. 2017;145: 1649–1657. doi:10.1017/S0950268817000358

183. Ospina J, Hincapie-Palacio D, Ochoa J, Molina A, Rúa G, Pájaro D, et al. Stratifying the potential local transmission of Zika in municipalities of Antioquia, Colombia. Trop Med Int Health TM IH. 2017;22: 1249–1265. doi:10.1111/tmi.12924

184. Reiner RC, Stoddard ST, Scott TW. Socially structured human movement shapes dengue transmission despite the diffusive effect of mosquito dispersal. Epidemics. 2014;6: 30–36. doi:10.1016/j.epidem.2013.12.003

185. Rojas D, Dean N, Yang Y, Kenah E, Quintero J, Tomasi S, et al. The epidemiology and transmissibility of Zika virus in Girardot and San Andres island, Colombia, September 2015 to January 2016. Euro Surveill. 2016;21. doi:10.2807/1560-7917.ES.2016.21.28.30283

186. Siraj AS, Oidtman RJ, Huber JH, Kraemer MUG, Brady OJ, Johansson MA, et al. Temperature modulates dengue virus epidemic growth rates through its effects on reproduction numbers and generation intervals. PLoS Negl Trop Dis. 2017;11: e0005797. doi:10.1371/journal.pntd.0005797

187. Yang HM, Boldrini JL, Fassoni AC, Freitas LFS, Gomez MC, de Lima KKB, et al. Fitting the Incidence Data from the City of Campinas, Brazil, Based on Dengue Transmission Modellings Considering Time-Dependent Entomological Parameters. PloS One. 2016;11: e0152186. doi:10.1371/journal.pone.0152186

188. Yang HM. The transovarial transmission in the dynamics of dengue infection: Epidemiological implications and thresholds. Math Biosci. 2017;286: 1–15. doi:10.1016/j.mbs.2017.01.006

189. Campbell KM, Haldeman K, Lehnig C, Munayco CV, Halsey ES, Laguna-Torres VA, et al. Weather Regulates Location, Timing, and Intensity of Dengue Virus Transmission between Humans and Mosquitoes. PLoS Negl Trop Dis. 2015;9: e0003957. doi:10.1371/journal.pntd.0003957

190. Masud MA, Kim BN, Kim Y. Optimal control problems of mosquito-borne disease subject to changes in feeding behavior of Aedes mosquitoes. Biosystems. 2017;156–157: 23–39. doi:10.1016/j.biosystems.2017.03.005

191. Lee J-S, Carabali M, Lim JK, Herrera VM, Park I-Y, Villar L, et al. Early warning signal for dengue outbreaks and identification of high risk areas for dengue fever in Colombia using climate and non-climate datasets. BMC Infect Dis. 2017;17: 480. doi:10.1186/s12879-017-2577-4

192. Ernst KC, Walker KR, Reyes-Castro P, Joy TK, Castro-Luque AL, Diaz-Caravantes RE, et al. Aedes aegypti (Diptera: Culicidae) Longevity and Differential Emergence of Dengue Fever in Two Cities in Sonora, Mexico. J Med Entomol. 2017;54: 204–211. doi:10.1093/jme/tjw141

193. Fitzgibbon WE, Morgan JJ, Webb GF. An outbreak vector-host epidemic model with spatial structure: the 2015-2016 Zika outbreak in Rio De Janeiro. Theor Biol Med Model. 2017;14: 7. doi:10.1186/s12976-017-0051-z

194. Velasquez-Castro J, Anzo-Hernandez A, Bonilla-Capilla B, Soto-Bajo M, Fraguela-Collar A. Vector-borne disease risk indexes in spatially structured populations. PLoS Negl Trop Dis. 2018;12. doi:10.1371/journal.pntd.0006234

195. Carbajo AE, Vezzani D. Waiting for chikungunya fever in Argentina: spatio-temporal risk maps. Mem Inst Oswaldo Cruz. 2015;110: 259–262. doi:10.1590/0074-02760150005

196. Manore CA, Hickmann KS, Xu S, Wearing HJ, Hyman JM. Comparing dengue and chikungunya emergence and endemic transmission in A. aegypti and A. albopictus. J Theor Biol. 2014;356: 174–191. doi:10.1016/j.jtbi.2014.04.033

197. Quiner CA, Parameswaran P, Ciota AT, Ehrbar DJ, Dodson BL, Schlesinger S, et al. Increased replicative fitness of a dengue virus 2 clade in native mosquitoes: potential contribution to a clade replacement event in Nicaragua. J Virol. 2014;88: 13125–13134. doi:10.1128/JVI.01822-14

198. Quintero-Gil DC, Ospina M, Osorio-Benitez JE, Martinez-Gutierrez M. Differential replication of dengue virus serotypes 2 and 3 in coinfections of C6/36 cells and Aedes aegypti mosquitoes. J Infect Dev Ctries. 2014;8: 876–884.

199. Serrato-Salas J, Izquierdo-Sánchez J, Argüello M, Conde R, Alvarado-Delgado A, Lanz-Mendoza H. Aedes aegypti antiviral adaptive response against DENV-2. Dev Comp Immunol. 2018;84: 28–36. doi:10.1016/j.dci.2018.01.022

200. Vazeille M, Gaborit P, Mousson L, Girod R, Failloux A-B. Competitive advantage of a dengue 4 virus when co-infecting the mosquito Aedes aegypti with a dengue 1 virus. BMC Infect Dis. 2016;16: 318. doi:10.1186/s12879-016-1666-0

201. Cromwell EA, Stoddard ST, Barker CM, Van Rie A, Messer WB, Meshnick SR, et al. The relationship between entomological indicators of Aedes aegypti abundance and dengue virus infection. PLoS Negl Trop Dis. 2017;11: e0005429. doi:10.1371/journal.pntd.0005429

202. Lana RM, Gomes MF da C, Lima TFM de, Honório NA, Codeço CT. The introduction of dengue follows transportation infrastructure changes in the state of Acre, Brazil: A network-based analysis. PLoS Negl Trop Dis. 2017;11: e0006070. doi:10.1371/journal.pntd.0006070

203. Eisen L, García-Rejón JE, Gómez-Carro S, Nájera Vázquez M del R, Keefe TJ, Beaty BJ, et al. Temporal correlations between mosquito-based dengue virus surveillance measures or indoor mosquito abundance and dengue case numbers in Mérida City, México. J Med Entomol. 2014;51: 885–890.

204. Liebman KA, Stoddard ST, Reiner RC, Perkins TA, Astete H, Sihuincha M, et al. Determinants of heterogeneous blood feeding patterns by Aedes aegypti in Iquitos, Peru. PLoS Negl Trop Dis. 2014;8: e2702. doi:10.1371/journal.pntd.0002702

205. Oliveira R de MAB, Araújo FM de C, Cavalcanti LP de G. Entomological and epidemiological aspects of dengue epidemics in Fortaleza, Ceará, Brazil, 2001-2012. Epidemiol E Serv Saude Rev Sist Unico Saude Bras. 2018;27: e201704414. doi:10.5123/s1679-49742018000100014

206. Monaghan AJ, Morin CW, Steinhoff DF, Wilhelmi O, Hayden M, Quattrochi DA, et al. On the Seasonal Occurrence and Abundance of the Zika Virus Vector Mosquito Aedes Aegypti in the Contiguous United States. PLoS Curr. 2016;8. doi:10.1371/currents.outbreaks.50dfc7f46798675fc63e7d7da563da76

207. Lo D, Park B. Modeling the spread of the Zika virus using topological data analysis. PloS One. 2018;13: e0192120. doi:10.1371/journal.pone.0192120

208. Stewart-Ibarra AM, Lowe R. Climate and non-climate drivers of dengue epidemics in southern coastal ecuador. Am J Trop Med Hyg. 2013;88: 971–981. doi:10.4269/ajtmh.12-0478

209. da Rocha Taranto MF, Pessanha JEM, dos Santos M, dos Santos Pereira Andrade AC, Camargos VN, Alves SN, et al. Dengue outbreaks in Divinopolis, south-eastern Brazil and the geographic and climatic distribution of Aedes albopictus and Aedes aegypti in 2011-2012. Trop Med Int Health TM IH. 2015;20: 77–88. doi:10.1111/tmi.12402

210. Souza-Neto JA, Powell JR, Bonizzoni M. Aedes aegypti vector competence studies: A review. Infect Genet Evol J Mol Epidemiol Evol Genet Infect Infect0. 2019;67: 191–209. doi:10.1016/j.meegid.2018.11.009

211. Lourenço-de-Oliveira R, Marques JT, Sreenu VB, Atyame Nten C, Aguiar ERGR, Varjak M, et al. Culex quinquefasciatus mosquitoes do not support replication of Zika virus. J Gen Virol. 2018;99: 258–264. doi:10.1099/jgv.0.000949

212. Tricco AC, Lillie E, Zarin W, O’Brien KK, Colquhoun H, Levac D, et al. PRISMA Extension for Scoping Reviews (PRISMA-ScR): Checklist and Explanation. Ann Intern Med. 2018;169: 467. doi:10.7326/M18-0850

